# Computationally-designed aptamers targeting RAD51-BRCA2 interaction inhibit RAD51 nuclear recruitment

**DOI:** 10.1101/2025.01.20.633558

**Authors:** Giulia Milordini, Elsa Zacco, Alexandros Armaos, Francesco Di Palma, Mirco Masi, Martina Gilodi, Jakob Rupert, Laura Broglia, Giulia Varignani, Michele Oneto, Marco Scotto, Roberto Marotta, Stefania Girotto, Andrea Cavalli, Gian Gaetano Tartaglia

## Abstract

The interaction between RAD51 and BRCA2 plays a key role in homologous recombination (HR), a critical DNA repair mechanism essential for the survival of cancer cells. Disrupting this interaction increases the sensitivity of cancer cells to chemotherapeutic agents. Here, we employed *in silico* methods to design a novel class of aptamers—customized single-stranded oligonucleotides—specifically engineered to bind RAD51. These aptamers were developed with the aim of selectively modulating RAD51’s nuclear recruitment and its role in DNA repair processes. The leading candidate displays high affinity for RAD51, competing with BRCA2 for the same interaction site *in vitro*, as confirmed through biolayer interferometry (BLI) and fluorescence lifetime imaging microscopy (FLIM). We tested the efficacy of the leading aptamer in pancreatic cancer cells and observed that it significantly impedes RAD51 nuclear localization, reduces homologous recombination (HR) efficiency, and increases DNA damage. Critically, our aptamer potentiates the cytotoxicity of the PARP inhibitor olaparib, exploiting synthetic lethality (SL) to induce cancer cell death. Our study showcases an aptamer-based approach for selectively targeting protein interactions within DNA repair pathways, introducing a promising avenue for SL-based treatments applicable to a wide range of cancers.

## INTRODUCTION

Cell growth and division entail the occurrence of DNA damage, which is usually answered by the activation of one or more specific DNA repair pathways^1^. Due to their fast rate of proliferation, in cancer cells there is a higher chance of damage accumulation and, consequently, a more frequent response to damage^2,3,4^. The more they grow, the better cancer cells become in efficiently activating DNA repair mechanisms^2,3^. The choice of the mechanism to be activated depends on the type of damage, which could be, for example, DNA strand breakages, base loss or deamination^5^.

In the case of double strand DNA break (DSB), the cellular repair primarily involves two pathways: homologous recombination (HR) and non-homologous end joining (NHEJ)^6^. The choice between these pathways is vital for maintaining genomic stability and cell viability and is dictated by the phase of the cell cycle and the specific characteristics of the DNA break^7^.

HR is a frequently adopted high-fidelity repair pathway that employs a homologous DNA sequence –usually the sister chromatid– as a template to accurately repair the DSB. Central to the HR pathway is the protein RAD51, involved in strand invasion and homology search^8,9^. RAD51 achieves these functions by forming helical filaments, whose structural properties depend on the interaction of the DNA to be repaired. RAD51’s structural adaptability enables it to play a pivotal role in HR by mediating strand invasion and homology search^10^.

Normally found in the cytosol, RAD51 is recruited to the site of DNA damage in the nucleus by direct association with BRCA2, a tumor suppressor protein. BRCA2 does not just passively recruit RAD51, instead it actively plays the role of a chaperone by ensuring that RAD51 is correctly positioned onto the single-stranded DNA regions that are exposed due to the damage. Once RAD51 is bound to these regions, it forms structures known as nucleoprotein filaments on the broken single-stranded DNA^11,12,13^. The filaments then search for and invade a homologous DNA duplex in a process called strand invasion, a critical step where the damaged DNA strand invades a sister chromatid to use it as a template for repair^8,14,15^. The physical interaction between RAD51 and BRCA2 is essential for the correct functioning of the HR pathway. For these reasons, targeting this complex has gained significant attention in drug discovery. In particular, BRC4, a 4-kDa peptide derived from one of the eight BRC repeats within BRCA2, has been exploited for its high-affinity binding to two specific pockets of RAD51 and has been selected to compete with the RAD51-BRCA2 interaction^16,17^. The first pocket, interacting with BRC4 motif FXXA, corresponds to the amino acids Ala157-Met210 on RAD51, while the second pocket, interacting with BRC4 motif LFDE, is formed by RAD51 residues Asn196-Ser214 and Ser239-Gly260^18^. Molecules that selectively target the RAD51-BRCA2 interface could potentially sensitize cancer cells to DNA-damaging agents, enhancing the efficacy of combined treatments^19,20^. The combination of anti-cancer drugs is a widely used way to improve therapeutic outcome leveraging on the principle of synthetic lethality (SL)^21^.

SL is the strategy by which two genes or pathways are disrupted at the same time, and such simultaneous intervention on two fronts leads to cell death. The disruption of only one of the pathways, instead, is damaging but not lethal^22^. For example, in the context of BRCA mutations causing deficiency in the HR repair mechanism, SL can be achieved by the use of poly (ADP-ribose) polymerase (PARP) inhibitors, such as olaparib^23^, that effectively prevent PARP enzyme from completing the repair process in the context of the base excision repair (BER) pathway^24^. The synergistic combination of a PARP inhibitor such as olaparib with HR inhibitors in patients lacking BRCA mutations, and therefore with intact HR repair mechanisms, sensitizes cancer cells to treatment irrespective of the genetic mutation pool ^25,26^. This dual-targeting strategy holds promise for enhancing treatment outcomes across diverse cancer types by exploiting common DNA repair pathways and SL principles. Such therapeutic approach represents a paradigm shift in cancer care, potentially transforming treatment strategies and improving patient outcomes^27,28^.

In this work the aim was to impair the HR function of cancer cells by altering the interaction between RAD51 and BRCA2 using aptamers, and to investigate its efficacy in the context of SL when combined with a PARP inhibitor. Aptamers are short, single-stranded oligonucleotides that can bind to specific targets with high affinity and specificity thanks to their tridimensional structures and to their tailor-made sequences^29–31^. Compared to conventional small molecules or antibodies, aptamers present multiple benefits. They can be synthesized with high yields and low costs and are highly accessible for chemical modifications that can improve their selectivity and stability^32^. They have a reduced potential to trigger an immune reaction and are therefore generally non-toxic to the body^33^. Furthermore, thanks to their compact nature and distinct structure, aptamers can effectively infiltrate target tissues that might be not easy to reach for bulkier entities^34^. Aptamers have been employed in various fields including therapeutic, diagnostic, and research-oriented applications. Their flexibility and malleability render them promising candidates for future biomedical innovations, especially in targeted drug delivery and personalized healthcare approaches^31,35^.

To date, the primary method for aptamer production is the Systematic Evolution of Ligands by Exponential Enrichment (SELEX) laboratory procedure, which involves repeated cycles of selection and amplification of the potential candidates^29,36^. Although it can generate aptamers with high affinity for the target protein, SELEX is still a time-consuming, labor-intensive technique, limited in its capacity to produce highly specific oligonucleotides. To address the need for speed and accuracy, we employed an *in silico* approach for the design of DNA aptamer to target the RAD51-BRCA2 interaction. The aptamers were developed using the in-house algorithm *cat*RAPID, a well-established computational tool to predict protein-nucleic acids interaction propensities by combining secondary structure, hydrogen bonding and van der Waals contributions^35,37,38^. The results presented here unveil novel sequences designed to selectively interfere with RAD51-BRCA2 interaction, aiming to disrupt the HR mechanism. Employing aptamers to this end could represent a solid starting point towards the development of nucleic acid-based drugs against multiple types of cancers.

## RESULTS

### *IN SILICO-*DESIGNED APTAMERS BIND A SPECIFIC REGION OF RAD51

We developed an *in silico* pipeline to design DNA aptamers that selectively bind to specific regions of RAD51 (**Material and Methods**). To interfere with RAD51 interactions with BRCA2, we focused on two key areas around the RAD51-BRC4 interaction pockets, FXXA and LFDE, which we named Region 1 and Region 2 (**Supplementary Fig. 1A and B**). Additionally, we included a third area, known as the helix-hairpin-helix (HhH) motif, involved in non-sequence-specific DNA binding. This part, designated as Region 3, serves as an internal control to evaluate whether the binding in Region 1 or Region 2 is stronger (**Supplementary Fig. 1C**). The *cat*RAPID algorithm was used to design aptamers of 9-16 nucleotides in length^39,40^. This falls within the range of 10–20 nucleotides that we previously exploited to successfully design aptamers^35^. Indeed, by limiting aptamers’ length, we aim to avoid complex structures, such as G-quadruplexes^41^, and ensure direct interaction with the binding interface. Longer structures can indeed expose additional sites for unintended protein interactions, increasing the risk of off-target effects^42,43^.

For all resulting sequences, an interaction score with each of the defined RAD51 regions was predicted, and the aptamers were ranked based on these scores, with the condition that the values were higher than those of a negative protein control, actin B (**Material and Methods**). The results consist in a series of aptamers characterized by repetitive short motifs rich in G. When comparing the *cat*RAPID scores relative to the predicted interaction propensities, all of the generated sequences display a higher value for Region 2 compared to Region 1 or Region. According to the predictions, the aptamers generated with *cat*RAPID would preferentially bind Region 2 of RAD51, specifically disturbing the interaction with BRC4 on the pocket LFDE. As a negative control, the interaction scores for rcApt –the reverse complementary sequences of the top scoring aptamer (Apt1)– was also included and the scores resulted to be close to 0.

### INTERACTION STUDIES VALIDATE THE PREDICTED BINDING OF SELECTED APTAMERS TOWARDS RAD51

The predicted binding propensities between the aptamers with an interaction score for Region 2 >8 and purified RAD51 were validated (**Supplementary Fig. 3**). The negative control rcApt was also tested. The binding affinities of the aptamers were determined by identifying their dissociation constants (K_d_s) using biolayer interferometry (BLI): the stronger the interaction, the lower the K_d_ (**Fig. 1**). To ensure a tight binding towards the target protein, the aim was to identify those aptamer sequences with a K_d_ for RAD51 <1 µM.

**Figure 1.**
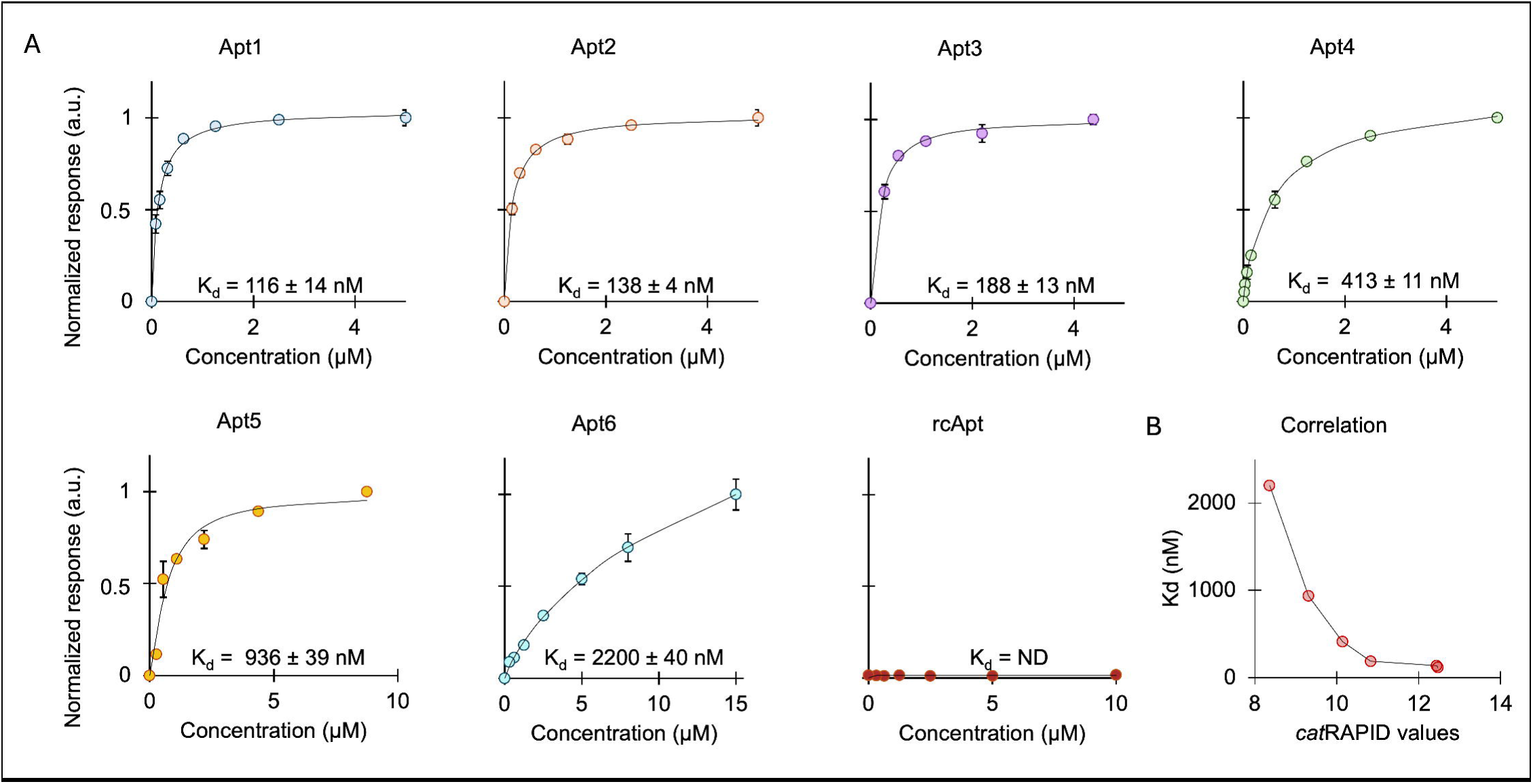
Analysis of aptamer binding to RAD51. **A**: BLI analysis illustrating the binding kinetics of the first 6 aptamers (Apt1 to Apt6) and a negative control (rcApt) with RAD51 protein at 25°C. Each curve represents the interaction profile of an individual aptamer, from which their respective binding affinities towards RAD51 can be extrapolated. Spheres: experimental points; line: curve fitting. The curve relative to rcApt is reported as raw data, since no binding response was registered. Three replicates were performed, and the mean and standard deviation were calculated for each point within the binding curves. Below each curve, the dissociation constant (K_d_) is indicated. (ND= not detectable); **B**: A scatter plot showing the correlation between experimentally determined K_d_s of the aptamers with RAD51 and their affinities predicted by the *cat*RAPID algorithm for Region 2.

The highest *cat*RAPID interaction scores, 12.47 and 12.44 respectively, were found for Apt1 and Apt2, which exhibited the lowest determined K_d_s of 116 ± 14 nM and 138 ± 4 nM (**Fig. 1A**). K_d_s of 188 ± 13 nM and 413 ± 11 nM were displayed by Apt3 and Apt4, both associated with a *cat*RAPID score of around 10. K_d_s of 936 ± 39 nM and exceeding 2 µM were presented by Apt 5 and 6, both assigned *cat*RAPID interaction scores below 9 (**Fig. 1A**). As predicted, the negative control rcApt exhibited no binding (**Fig. 1A**).

The results of these analyses indicate that the sequences of Apt1-Apt6, predicted by *cat*RAPID to interact strongly with RAD51, displayed a trend where a higher *cat*RAPID score correlated with lower K_d_ values (**Fig. 1A**). A notably robust negative correlation of −0.85 was uncovered, thereby confirming that the affinity trend is in accordance with the *cat*RAPID score and that the algorithm was able to correctly predict the aptamers with the highest binding probability (**Fig. 1B**).

### AN *IN SILICO*-DESIGNED APTAMER CAN DISRUPT RAD51 FILAMENTS *IN VITRO*

Following the prediction of binding propensities between the top 6 aptamer candidates and RAD51, their effect on the protein oligomerization state was investigated. Apt1 was selected as the aptamer with the highest affinity based on computational predictions and *in vitro* evaluations.

The impact of Apt1 on RAD51 was assessed using negative staining transmission electron microscopy (TEM), incubating RAD51 with increasing concentrations of Apt1 (**Fig. 2** and **Supplementary Fig. 4A-B**). In agreement with the literature, also in this study RAD51 in isolation forms mainly worm-like helical fibrils (**Fig. 2A**)^44^. Upon incubation of RAD51 with equal molar concentrations of Apt1, no visible effect was observed (**Fig. 2B**). However, increasing the concentration of Apt1 to six times that of RAD51 resulted in the almost total disruption of the long worm-like fibrils that were replaced by smaller oligomers (**Fig. 2C**). The resulting ring-like arrangements of the oligomers displayed an average diameter of 13.3 ± 1.4 nm (**Supplementary Table 1**). A further increase in the concentration of Apt1 to twelve times that of RAD51 led to the complete disruption of the oligomers (**Fig. 2D**), indicating that the aptamer significantly affected RAD51 quaternary structure, particularly in terms of filament stability. Incubating RAD51 with the highest concentration of the negative control aptamer, rcApt, did not perturb the filamentous structure of the protein (**Supplementary Fig. 4C**).

**Figure 2.**
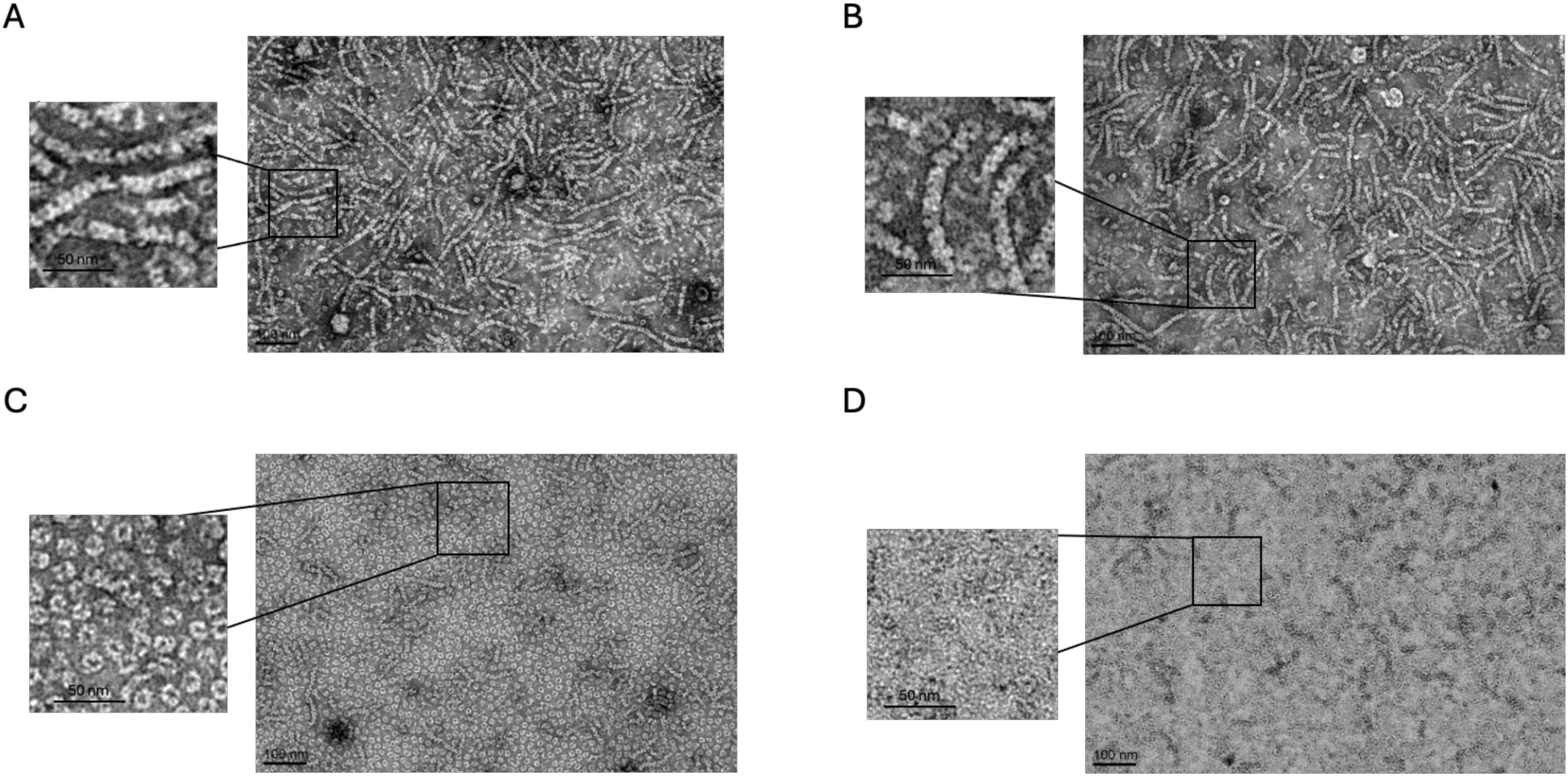
Negative staining TEM images of RAD51 fibrils/oligomers in the presence and in the absence of Apt1. **A:** RAD51 by itself, displaying the characteristic fibril structure; **B:** RAD51 incubated with Apt1 at a molar ratio of 1:1, exhibiting similar supramolecular structures; **C:** RAD51 incubated with Apt1 at a molar ratio of 1:6, revealing a significant reduction in the presence of long fibrils, accompanied by the formation of smaller oligomeric structures; **D.** RAD51 incubated with Apt1 at a molar ratio of 1:12, resulting in the complete abrogation of fibrils.

These results indicate an interference exerted selectively by Apt1 —and not by rcApt— on the oligomerization state of RAD51. By interfering with RAD51 quaternary structure, the aptamer might also affect the protein function to repair the damaged DNA via HR.

### COMPUTATIONAL PREDICTIONS CONFIRMS THE STRUCTURAL ALTERATIONS IN RAD51 FILAMENTS INDUCED BY APT1

To gain atomistic details on the mechanisms by which Apt1 disassembles RAD51 filaments, structural predictions of RAD51 fibrils in the presence of increasing copy number of Apt1 (1, 6 or 12) were performed using AlphaFold 3^45^ (**Fig. 3**). As a control, the same predictions were repeated with rcApt (**Supplementary Fig. 5**).

**Figure 3.**
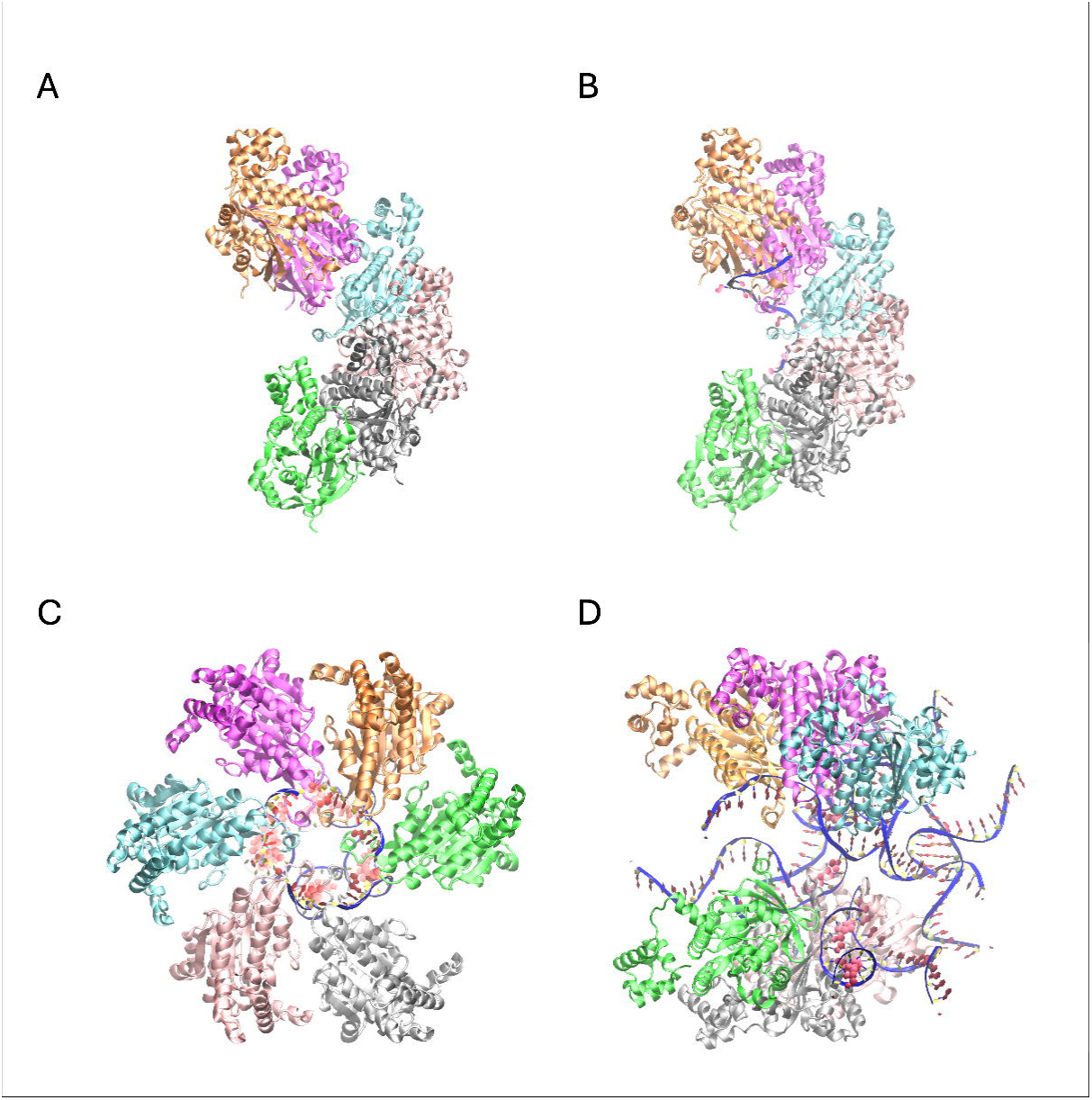
Structural prediction of RAD51 fibrils/oligomers in the presence and absence of Apt1. **A**: RAD51 by itself (6 protomers), displaying the characteristic fibril structure; **B**: One filament of RAD51 in complex with one Apt1 molecule, exhibiting a similar supramolecular structure with A; **C**: One filament of RAD51 in complex with six Apt1 molecules, revealing a reorganization of the protofilament into a hexameric annular structure; **D**: One filament of RAD51 in complex with 12 Apt1 molecules, resulting in the complete abrogation of the fibrils. In this latest case, two trimers are interacting with several copies of the aptamer. Each protein chain is shown in a different color using the cartoon-based representation (chain A in orange, B in magenta, C in cyan, D in pink, E in silver and F in green). Each aptamer copy is reported as a blue ribbon, with the sugar rings in yellow and the nucleobases in red.

Following up on one of the resolved structures of a RAD51 filament in which one helical turn corresponds to 6.6-7.3 RAD51 protomers per turn^46^, the prediction of the RAD51 filament was attempted with both six and seven monomers. It was found that the former was better suited to obtain a model (**Fig. 3A**) that closely resembled the X-ray helical structure of RAD51 in the absence of the ssDNA filament to be repaired, defined as *apo* protein (PDB ID: 5NWL)^46^. A relevant difference between the deposited structure and the model here proposed is the reconstruction of the unresolved loop from Ala271 to Gly288. This portion (part of L2 in Pellegrini et. al.^47^ connecting β-strand B5 to β-strand B6) was crucial to properly model the interaction with the aptamers^48^. The prediction of the structure including 7 RAD51 copies was perfectly overlapping with the one containing 6 copies, with the seventh protomer beginning a new helical turn, and it is not reported to avoid redundancy.

The model here proposed is characterized by a pitch of 125 Å, fitting the previously reported range of 112–134 Å^46^ for the *apo* protein. This reconstruction of RAD51 fibril structure was used as reference to evaluate the predicted complexes formed by the protein filament with increasing copies of Apt1 molecules.

In the model with one copy of Apt1, the aptamer interacts with five of the six RAD51 molecules forming the filament, without impacting its structure (**Fig. 3B**). In accordance with the design, the aptamer is located in a different groove (**Supplementary Fig. 2B**) compared to the protein’s ssDNA binding site identified by cryoEM (PDB ID: 8PBC)^49^. In particular, the aptamer lies juxtaposed to the DNA binding site used for repair, sharing with it the interaction with the backbone atoms of few amino acids only (**Supplementary Table 2**). In this configuration, the pitch length shrunk to 115 Å, still in accordance with the deposited *apo* structure (second assembly of the 5NWL PDB)^46^. In this framework, 17 out of 25 predicted models were found binding Apt1 at a different spot than the RAD51 ssDNA binding site (**Fig. 3B**); in other 5 configurations, Apt1 was bound to the ssDNA binding site; in 3 out of 25 models, Apt1 was binding the filament at different spots with weak and non-specific interaction, or not binding at all.

To interpret the TEM results, the effect of 6 copies of Apt1 on the disassembly of one RAD51 filament was predicted (**Fig. 3C**). The resulting model revealed the loss of the helical turn of RAD51 filament and the rearrangement of the six RAD51 protomers into a circular complex, with each of the aptamer molecules intercalating between adjacent chains. This peculiar structure for a RAD51 oligomer displayed an approximate diameter of 13 nm, qualitatively resembling the smaller non-filamentary oligomeric structures identified by TEM imaging at the 1:6 molar ratio (**Fig. 3C**). According to the structural predictions, the amino acids of RAD51 filament involved in the interaction with the aptamer only partially overlapped with the ones found in the model comprising one RAD51 filament and one Apt1 molecule (**Supplementary Table 2**): among them, Arg229, Arg235, Leu238, Ser239, Ala271 and Asn290. Conversely, the unique intermolecular interactions borne by the annular shaped complex involved some of the residues of the reconstructed loop: Phe279, Ala280, Ala281, Asp282 and Pro283. Statistically, the proposed circular oligomeric structure was observed in 15 out of 25 predictions. Conversely, in only 5 out of 25 predictions, the RAD51 filament was able to maintain its structure in the presence of six copies of Apt1. Among these, five showed non-specific interactions with the filament, and only one bound to the ssDNA binding site. In the remaining 5 predictions, the oligomeric structure was disrupted, yielding a mix of dimers and trimers connected by aptamers.

Recapitulating the TEM condition with the highest amount of aptamer, RAD51 filament was then modeled with 12 Apt1 molecules. None of the AlphaFold 3 generated structures resulted in a helical filament, nor as a circular structure. Instead, the putative filament is broken in two parts, resulting in two trimers interconnected with the DNA aptamers, mostly in a non-specific arrangement (**Fig. 3D**). Indeed, nine out of 12 aptamer strands were poorly bound to the proteins by means of non-specific and non-conserved contacts, lying between the two trimers and establishing occasional interchain base pairs with other aptamer copies. There are two exceptions: i) two Apt1 molecules binding the RAD51 protomers (**Fig. 3D**) at a similar spot as in the one filament-one aptamer model (**Fig. 3B**), mostly via the same amino acids (**Supplementary Table 2**), albeit with a different binding mode; ii) another aptamer contacting the protein between chain E and F (in silver and green, respectively, in **Fig. 3D**), close to Asp274-Pro283.

Both the identified putative protein patches involved in the interaction with the nucleotides of Apt1 are part of the Region 2 of RAD51, the target area for which the aptamers were specifically designed (**Material and Methods**, **Supplementary Fig. 1B and 2** and **Supplementary Table 2**). Instead, in contrast with Apt1, the structural predictions in the presence of an increasing copy number of rcApt show that also *in silico* the control DNA aptamer is not able to interfere with the RAD51 filament formation (**Supplementary Fig. 5**).

### THE APTAMER AFFECTS BRC4-RAD51 INTERACTION

To rigorously assess the binding specificity of the designed aptamers to the targeted pockets on RAD51, Fluorescence Lifetime Imaging Microscopy (FLIM) was employed. This technique enables us to probe molecular interactions with high precision by measuring changes in fluorescence lifetimes, which can indicate binding events at a molecular level^50,51^. Here, FLIM was used to investigate whether the aptamers could effectively influence the interaction between RAD51 and the BRC4 peptide.

The goal of this experiment was to determine whether the aptamers could successfully affect BRC4 binding on RAD51. By targeting this specific area, the aptamers could effectively inhibit its interaction with BRCA2. As a proof of concept, the best performing aptamer, Apt1, was selected to be studied with this technique and labeled with Texas Red (**Fig. 4**). As a control, rcApt, which does not bind RAD51 (**Fig. 4**), was included.

**Figure 4.**
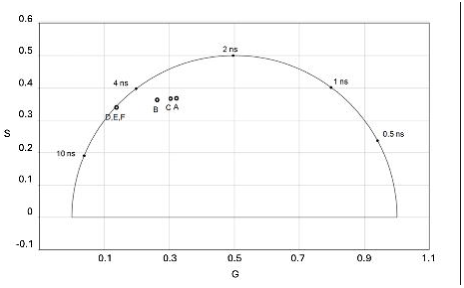
FLIM analysis of Apt1 affecting RAD51 and BRC4 interaction. Phasor plot from FLIM analysis showing sine component (S) and cosine component (G) on the axes, for the fluorophore Texas Red conjugated to the aptamers Apt1 and rcApt. Points **A** to **F** correspond to different experimental conditions. **A**: Apt1 alone; **B**: Apt1 after incubation with RAD51; **C**: Apt1 with RAD51 followed by the addition of BRC4; **D**: rcApt alone; **E**: rcApt after incubation with RAD51; **F**: rcApt with RAD51 followed by the addition of BRC4. **D**-**F** results are perfectly overlapping.

The FLIM experiments were performed in various conditions: the lifetime fluorescence of Texas Red directly conjugated to Apt1 and conjugated to rcApt was characterized by themselves, in the presence of RAD51, and with RAD51 together with BRC4. The fluorescence lifetime decay of Apt1 was analyzed before (**Fig. 4**, point **A**) and after (**Fig. 4**, point **B**) the addition of RAD51, revealing an increase in the slow component (τ_slow_) from 5.01 ± 0.1 ns to 5.32 ± 0.2 ns (**Table 1**). This increase of τ_slow_ indicates an interaction between RAD51 and Apt1. Introducing BRC4 into this system (**Fig. 4**, point **C**) led to a reduction in τ_slow_ down to 5.09 ns, suggesting a partial displacement of Apt1 from its binding site by BRC4, thus reintroducing the aptamer into solution. The observed intermediate lifetime of 5.09 ± 0.1 ns supports the theory that Apt1 was both in solution and partially bound to RAD51, indicating that its interaction with RAD51 affects the protein’s capacity to bind BRCA2.

**Table 1.** The table reports the faster (τ**_fast_**) and slower (τ**_slow_**) components of the fluorescence decay lifetime, sine component (S), cosine component (G), and values for fast and slow components ratio obtained from fluorescence lifetime imaging microscopy (FLIM). These measurements refer to the fluorophore Texas Red conjugated to Apt1 and rcApt aptamers, detailing variations upon the addition of RAD51 and BRC4. The observed change in **t** values for Apt1 in the presence of RAD51 substantiates a binding between them. On the other hand, the **t** values for the control molecule, rcApt, remain unchanged, confirming its lack of interaction with RAD51.

| Point | Condition | $\tau_{\text{fast}}$ (ns) | $\tau_{\text{slow}}$ (ns) | G (°) | S (°) | Fast component ratio | Slow component ratio |
| --- | --- | --- | --- | --- | --- | --- | --- |
| A | Apt 1 | $1.11 \pm 0.2$ | $5.01 \pm 0.1$ | 0.322 | 0.368 | 0.29 | 0.71 |
| B | Apt1 + RAD51 | $1.34 \pm 0.1$ | $5.32 \pm 0.2$ | 0.226 | 0.355 | 0.18 | 0.82 |
| C | Apt1 + RAD51 + BRC4 | $1.22 \pm 0.3$ | $5.09 \pm 0.1$ | 0.304 | 0.367 | 0.29 | 0.71 |
| D | rcApt | - | $5.1 \pm 0.1$ | 0.128 | 0.329 | - | 1.0 |
| E | rcApt + RAD51 | - | $5.1 \pm 0.1$ | 0.128 | 0.329 | - | 1.0 |
| F | rcApt + RAD51 + BRC4 | - | $5.1 \pm 0.1$ | 0.128 | 0.329 | - | 1.0 |

Furthermore, the decomposition of the fluorescence signal into ratios of “slow” and “fast” components revealed the presence of multiple fluorescent states, each differently influencing the fluorescence lifetime. The fast component, indicating a short lifetime, suggests rapid return to the ground state, whereas the slow component, indicating a longer lifetime, suggests a slower return. The increase in the Apt1’s slow component from 0.71 to 0.82 upon adding RAD51, and its subsequent return to 0.71 with the addition of BRC4, supports that BRC4 and Apt1 both target a common region of RAD51 (**Table 1**). Predicted complexes from AlphaFold 3 showing the positioning of BRC4 and Apt1 at their respective binding sites on the RAD51 filament are presented in **Supplementary Fig. 6,** along with the model of the RAD51 oligomer in the presence of excess BRC4 peptide (**Supplementary Fig. 7**). A parallel experiment with rcApt consistently showed a constant τ_slow_ of 5.01 ± 0.1 ns under all tested conditions: by itself, with RAD51, and with RAD51 plus BRC4 (**Fig. 4**, points **D**, **E**, **F**). This uniformity in decay time irrespective of the presence of RAD51 or BRC4 across these conditions confirms that the negative control rcApt does not interact with either molecules.

These findings provide strong evidence that the designed aptamers are interacting with the intended regions on RAD51, validating the *in silico* predictions and confirm that the aptamers have the potential to disrupt the RAD51-BRCA2 interaction, offering a promising strategy for impairing the homologous recombination pathway in cancer cells.

### THE APTAMER ALTERS THE CELLULAR LOCALIZATION OF RAD51, IMPAIRING THE FORMATION OF RAD51-POSITIVE *FOCI*

The performance of the aptamers against RAD51 activity in a cellular environment was assessed using BxPC-3 cells, derived from a human pancreatic adenocarcinoma^52^. These cells exhibit tumorigenicity, invasiveness, and drug resistance that mirror, in the limit of a 2D culture, the behavior of pancreatic cancer cells *in vivo*. The aptamer sequences were stabilized against nuclease activity with the introduction of locked bases. The fluorophore Texas red was conjugated to one end to enable their visualization with fluorescence microscopy techniques (**Material and Methods**). Apt1, the best performing aptamer, and its negative control rcApt were selected for further investigations.

Under physiological conditions, RAD51 is distributed mostly in the cytoplasm. To participate in DNA repair via HR, RAD51 is recruited from the cytosol into the nucleus and brought to the damage site by BRCA2^53,54^. Upon recruitment to the nucleus in response to DSBs, RAD51 accumulates at the damage site forming distinctive nuclear *foci,* regions of DNA where the damage occurs^55^.

One way to investigate the potential effect of the aptamers on the function of RAD51 is to evaluate the subcellular localization of the protein under stress and quantify the number of cells positive to RAD51 *foci*. To recreate stressed conditions, cisplatin was employed. Cisplatin is a double DNA breakage compound that forms covalent bonds with DNA bases, leading to the creation of adducts which can distort DNA’s structure^56,57^.

Immunofluorescence (IF) microscopy was employed to determine the effect of Apt1 and of the negative control rcApt on the subcellular distribution of diffused RAD51 in the presence and in the absence of cisplatin^58,59^ (**Fig. 5A** and **5B**). As expected, the nuclear fluorescence of RAD51 in control cells was significantly lower than post cisplatin-treated cells (27.9 ઱4.6 % vs 73.6 ±12.2 %) (**Fig. 5A** and **B**, top panels; **Fig. 5C**). Cells stressed with cisplatin and, at the same time, transfected with Apt1 showed a nuclear migration of RAD51 comparable to non-treated control cells (31.4 ±7.5 % fluorescence) (**Fig. 5A** and **B**, middle panels; **Fig. 5C**). Cells transfected with the DNA aptamer control rcApt mirrored the pattern of the cisplatin-treated cells (RAD51 nuclear fluorescence 73.80 ±12.7 %), indicating a non-significant impact on RAD51 nuclear recruitment (**Fig. 5A** and **B**, bottom panels; **Fig. 5C**).

**Figure 5.**
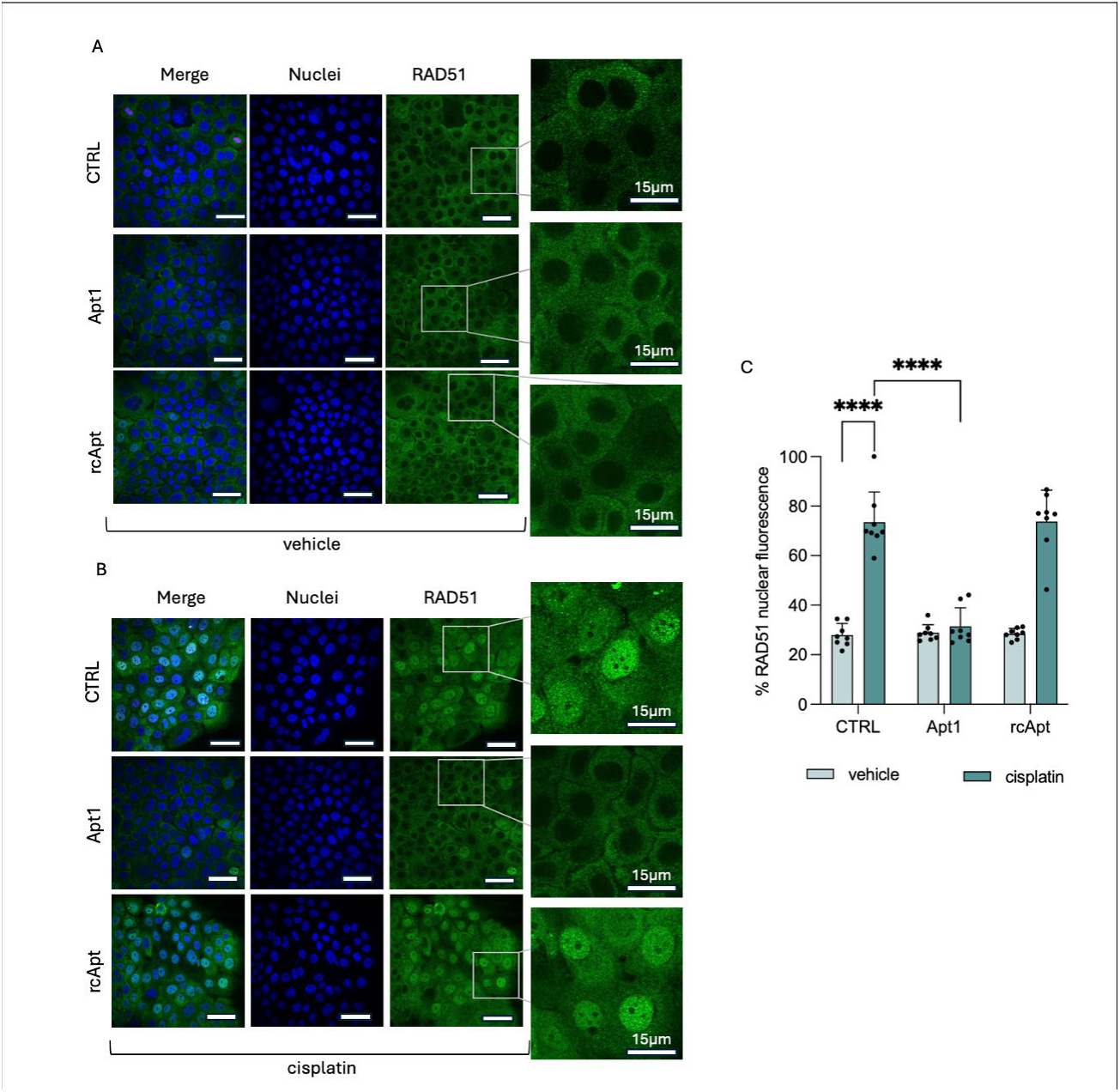
Impact of Apt1 on RAD51 cellular localization in response to DNA damage. **A**: IF images of non-stressed BxPC-3 cells, untreated (top panels), treated with Apt1 (middle panel), and treated with the negative control rcApt (bottom panels); **B**: IF images of cisplatin-stressed BxPC-3 cells, untreated (top panels), treated with Apt1 (middle panel), and treated with the negative control rcApt (bottom panels). Blue: DAPI-stained nuclei; green: immunostained RAD51. The excerpts on the right offer a magnified view of the cells; **C**: Bar chart illustrating RAD51 % nuclear fluorescence in physiological conditions (light green bars) and under cisplatin-induced stress (dark green bars). When not otherwise specified, scale bars measure 50 µm. Experiments were performed in biological triplicates, and data were acquired from eight images per sample for each condition (****= p-value <0.0001, calculated with Tukey test).

These results strongly suggest that, under stress-induced conditions, Apt1 markedly reduced RAD51’s nuclear transport.

IF microscopy was employed also to assess nuclear RAD51 *foci* formation in both physiological conditions and in stress-induced conditions (cisplatin). To quantify the RAD51-positive *foci*, the percentage of cells positive for *foci* detected in the presence of cisplatin was considered to be 100; under physiological conditions, RAD51 *foci* comprise approximately 10-20% of total *foci* (**Fig. 6A-B, top panels**). The addition of either Apt1 or rcApt did not vary this result (**Fig. 6A, middle and bottom panels**; **Fig. 6C**). When cells were stressed with cisplatin, the presence of Apt1 resulted in a significant reduction of ca. 75% in RAD51 *foci*, (**Fig. 6B, middle panels; Fig. 6C)**. The negative control rcApt did not have the same impact (**Fig. 6B, bottom panels; Fig. 6C**).

**Figure 6.**
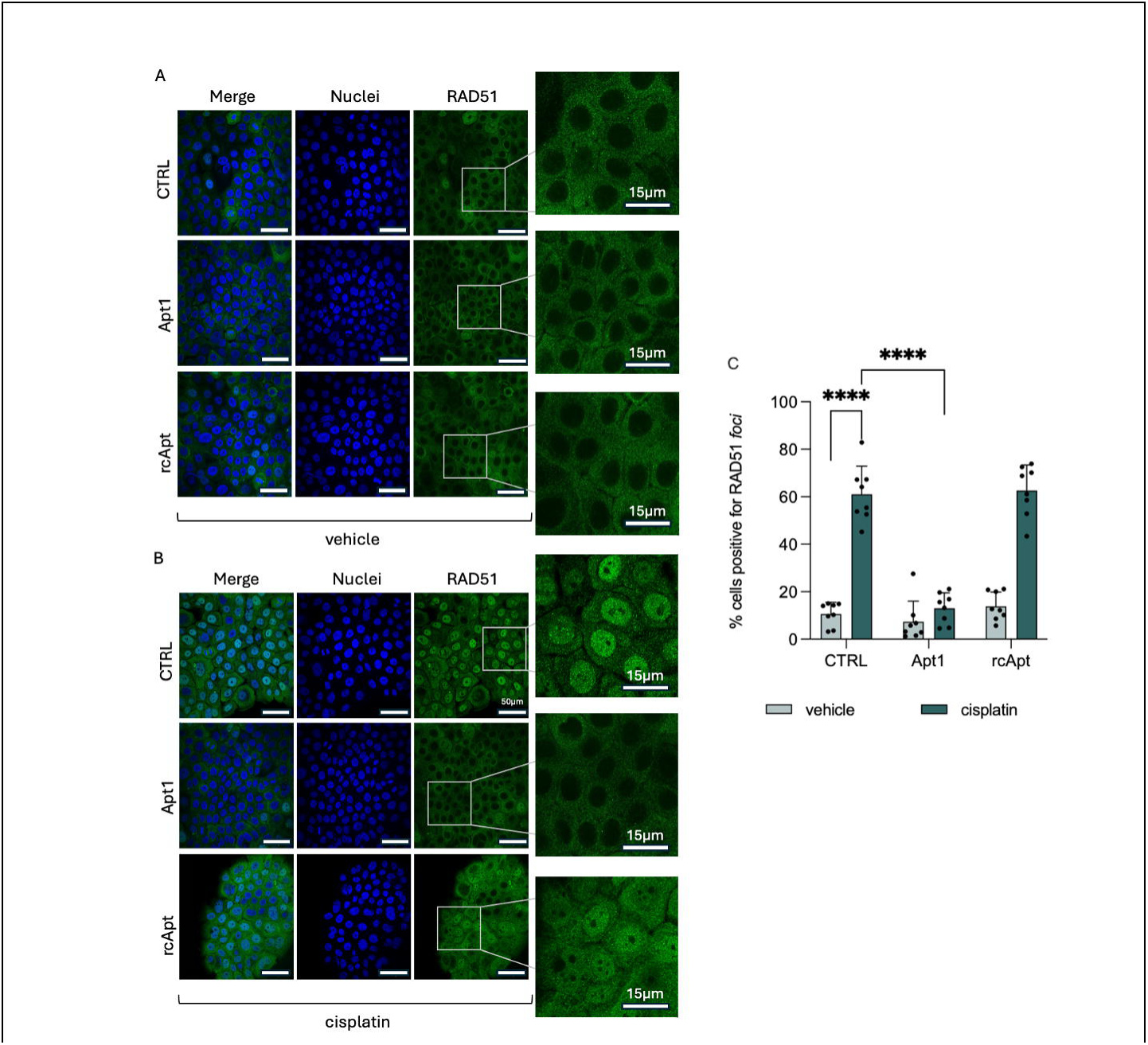
Impact of Apt1 on RAD51-positive *foci* formation in response to DNA damage. **A**: IF images of non-stressed BxPC-3 cells, untreated (top panels), treated with Apt1 (middle panel), and treated with the negative control rcApt (bottom panels); **B**: IF images of cisplatin-stressed BxPC-3 cells, untreated (top panels), treated with Apt1 (middle panel), and treated with the negative control rcApt (bottom panels). Blue: DAPI-stained nuclei; green: immunostained RAD51. The excerpts on the right offer a magnified view of the cells. **C**: Bar chart illustrating RAD51 nuclear *foci* levels in physiological conditions (light green bars) and under cisplatin-induced stress (dark green bars), highlighting Apt1’s role in diminishing RAD51’s nuclear *foci* number, a key factor in HR pathway activation for DNA repair. When not otherwise specified, scale bars measure 50 µm. Experiments were performed in biological triplicates, and data were acquired from eight images per sample for each condition (****= p-value <0.0001, calculated with Tukey test.

These results suggest a significant effect of Apt1 on the ability of RAD51 to form nuclear *foci*. The aptamer appears to have a negative impact on the function of RAD51 as a DNA repair protein.

Taken together, these findings emphasize the therapeutic promise of Apt1 in targeting DNA repair pathways and warrant further exploration in cancer treatment strategies.

### THE APTAMER PROMOTES DNA DAMAGE ACCUMULATION, LEADING TO CELL DEATH

Compared to normal cells, cancer cells are characterized by higher replication rates and overreliance on HR-mediated DNA damage repair^60^. These pathophysiologic conditions, coupled to DNA quality control deficiencies (e.g., due to *TP53* mutations) often observed in different tumour types, make cancer cells more susceptible to DNA damage accumulation^61–64^. In this regard, a prolonged HR inhibition results in increased DNA damage and genomic instability, effects that are further exacerbated upon the inhibition of PARP-mediated SSBs DNA repair mechanisms^3^. Therefore, to investigate the potential of the aptamers in halting DNA repair and amplifying olaparib-induced DNA damage accumulation, we analyzed DSB events by quantifying the number of *foci* formed by phosphorylated histone variant γH2AX^65^ using IF^65^. The baseline levels of DSBs were first investigated in cells left to grow for three days; this growth period enabled us to quantify both the presence of γH2AX *foci* and to evaluate the effect of our aptamers on their formation (**Fig. 7**). In the absence of aptamers, approximately 12% of all cells were positive for γH2AX *foci* (**Fig. 7A top panels and Fig. 7B**). However, in the presence of Apt1 a significant increase in DSBs was observed (**Fig. 7A middle panels and excerpt, and Fig. 7B**): the cells positive to *foci* reached 21.1%, almost double the percentage observed in both untreated controls and cells treated with the negative control rcApt, which was 10.9% (**Fig. 7A bottom panels and Fig. 7B**). This increase in DSBs suggested that Apt1 but not rcApt interfered with physiological repair processes, hinting at a potential role of Apt1 in modulating the DNA repair processes.

**Figure 7.**
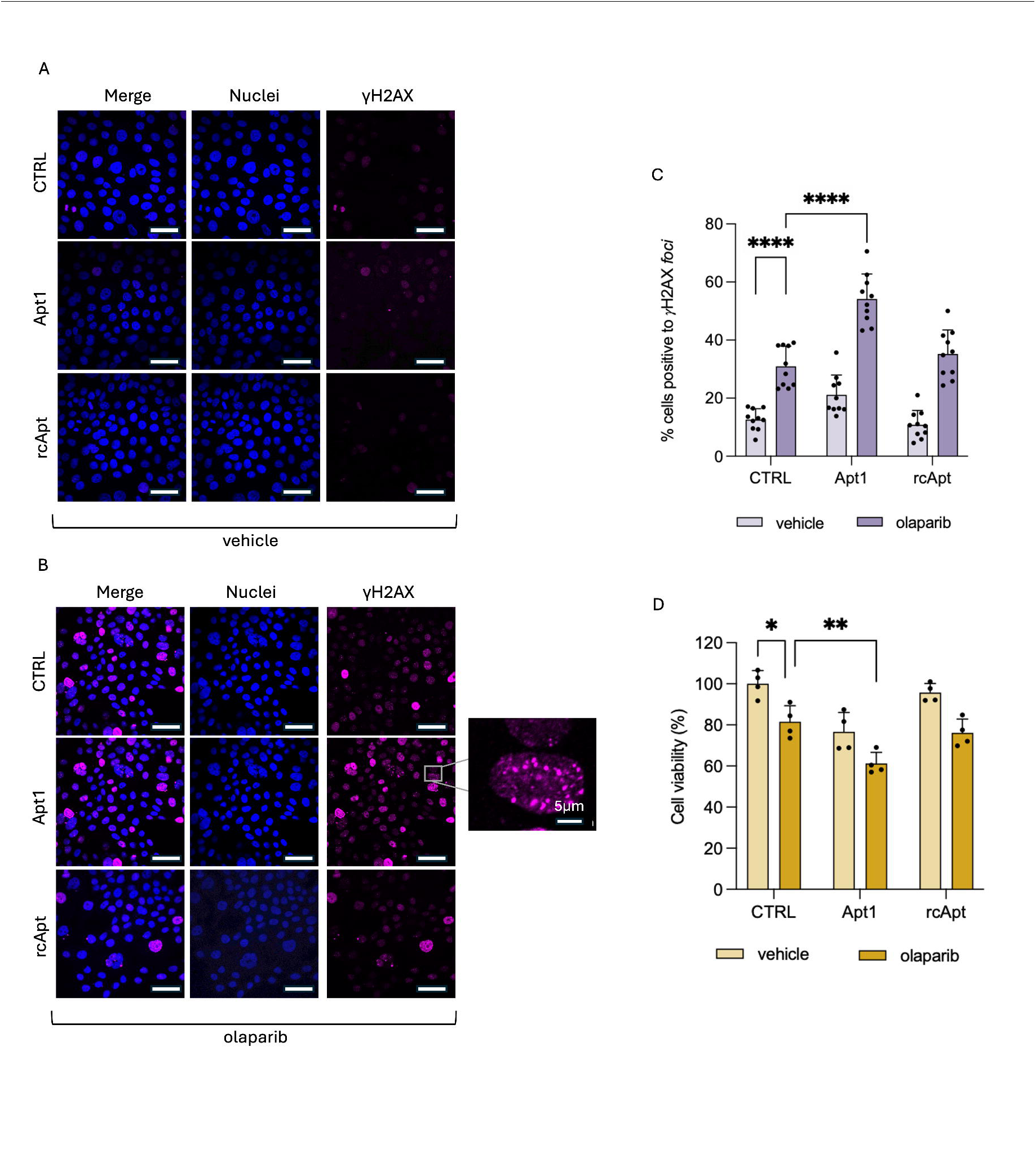
Impact of Apt1 on DNA repair mechanisms and on BxPC-3 cell viability in the context of SL. **A**: IF images of BxPC-3 cells positive for γH2AX *foci* in the absence of olaparib, 72 hours after aptamer transfection, untreated (**top panels**), treated with Apt1 (**middle panels**), and treated with the negative control rcApt (**bottom panels**); **B**: IF images of BxPC-3 cells positive for γH2AX *foci* 72 hours after aptamer transfection in the presence of olaparib, untreated (**top panels**), treated with Apt1 (**middle panels**), and treated with the negative control rcApt (**bottom panels**). The excerpt shows a cell positive to γH2AX *foci*; **C**: Bar chart illustrating the percentage of BxPC-3 cells positive for γH2AX *foci* 72 hours after aptamer transfection in the presence and in the absence of olaparib treatment. The comparison includes untreated cells (CTRL), cells treated with Apt1, and cells exposed to rcApt (negative control); **D**: Bar chart illustrating the percentage of viable BxPC-3 cells 96 hours post-aptamer transfection. The comparison includes untreated cells (CTRL) and cells treated with Apt1 and rcApt (negative control), in the presence and in the absence of olaparib treatment. (*= *p*-value <0.05)(**= p-value <0.01). When not otherwise specified, scale bars measure 50 µm. Experiments were performed in biological triplicates, and data were acquired from 10 images per sample for each condition (****= p-value <0.0001, calculated with Tukey test).

IF microscopy was employed to assess the effect of Apt1 and of the negative control rcApt on γH2AX levels, in the presence of the PARP inhibitor olaparib. Untreated cells exhibited a low percentage of cells positive for γH2AX *foci* (approximately 14%) (**Supplementary Fig. 7**). Cells treated with olaparib and Apt1 showed a significant increase in γH2AX *foci* (54.2%) compared to cells treated solely with olaparib (30.9%) (**Fig. 7B**). In contrast, cells treated with olaparib and rcApt showed a modest increase in γH2AX *foci* (35.3%) (**Fig. 7B**).

The increment in the number of these species is indicative of accumulation of DNA damage that escapes repair mechanisms. This result underscores the potent DNA-damaging effect induced by Apt1 in combination with olaparib.

Once established the effect of the aptamers on physiological accumulation of γH2AX *foci*, their impact on cell viability was investigated to confirm SL manifestation. Olaparib, a PARP inhibitor, is primarily used in cancer patients affected by mutations in the *BRCA1* or *BRCA2* genes^23,24^. These mutations impair the cells’ ability to repair DNA through the HR pathway, leading to increased reliance on alternative repair mechanisms, such as PARP-mediated base excision repair. By inhibiting PARP enzymes, olaparib exacerbates DNA damage accumulation in these already compromised cancer cells, ultimately leading to cell death. By combining olaparib with an aptamer acting as HR inhibitor, the aim is to sensitize target cells to treatment, regardless of their genetic pool.

To quantify the actual SL potential of Apt1 in combination with olaparib, viability tests were performed at the end of the treatment. The viability of BxPC-3 cells was assessed using the 3-(4,5-dimethylthiazol-2-yl)-2,5-diphenyltetrazolium bromide (MTT) assay, a widely employed method to evaluate cell metabolic activity and proliferation.

Viability was assessed in cells transfected with aptamers under both olaparib-treated and untreated conditions. In the absence of olaparib, control cells showed standard levels of viability, normalized to 100%, whereas cells treated with Apt1 exhibited a viability of 76%, underlying the negative effect of the aptamer on cell health (**Fig. 7D**). This reduction in viability was not observed with the negative control rcApt, where cells displayed a viability of approximately 95% (**Fig. 7D**).

In the presence of olaparib alone, viability decreased to 81% compared to untreated controls (**Fig. 7D**). When Apt1 was transfected in olaparib-treated cells, the viability further decreased to 61%, whereas cells treated with rcApt showed a viability compared to olaparib alone (76%) (**Fig. 7D**).

It is evident that the combination of olaparib and Apt1 resulted in a more significant decrease in cell viability, suggesting that Apt1 may enhance the cytotoxicity of olaparib, potentially through mechanisms that impair HR or promote cell death pathways. These findings highlight Apt1’s potential as a therapeutic agent, offering a promising strategy to exploit vulnerabilities in DNA repair pathways for improved cancer treatment outcomes.

## DISCUSSION

This study investigates the design of *in silico* aptamers that target specific pockets of the DNA-repair protein RAD51, interfering with its function and limiting its role in the cell’s DNA damage response. The focus is on altering the interaction between RAD51 and its recruitment protein BRCA2, which facilitates RAD51’s nuclear translocation and oligomerization at DNA damage sites. The recruitment of RAD51 and other repair proteins is crucial for the formation of nuclear *foci*, key hubs for DNA repair.

The interaction between RAD51 and BRCA2 has recently attracted the interest of the cancer community as a potential target for therapeutic intervention. Similarly to other studies, Bagnolini *et al.* used dynamic combinatorial chemistry to identify small molecules inhibiting this interaction^17,18^. Among these inhibitors, the most potent ones disrupted RAD51 *foci* formation and heightened cancer cell sensitivity to PARP inhibitors, which interfere with the DNA repair pathway called base excision repair. Scott *et al.* developed CAM833, a small molecule that inhibits the BRCA2-RAD51 interaction by disrupting RAD51 nucleoprotein filament formation and enhancing cell sensitivity to DNA damaging agents^18^. Moreover, Schipani *et al.* identified a peptide highly effective in targeting RAD51, shedding light on how BRCA2 mediates the defibrillation of RAD51 filaments, a critical step in HR^44^. These studies highlight the therapeutic potential of targeting RAD51-BRCA2 interaction to impair DNA repair and sensitize cancer cells to DNA damaging drugs.

In addition to small molecules and peptides, aptamers have emerged as inhibitors of RAD51, as demonstrated by Martinez *et al*. Three aptamers inhibiting the formation of the complex formed by RAD51 and ssDNA were identified through SELEX, although with unsatisfactory complex dissociation ability^66^. In oncology, aptamers are currently being used also to target other proteins implicated in cancer progression. For instance, AS1411^67,68^ and AGRO100^69,70^ target nucleolin, a protein over-expressed in cytoplasmic membranes of cancer cells, and exploit the selective binding of the aptamers to this protein for the targeted delivery of drugs conjugated to the aptamers. Another example is offered by NOX-A12, an aptamer targeting the chemokine CXCL12 that competes with glycosaminoglycans for the presentation of CXCL12 on the cell surface, leading to the chemokine neutralization^71^. These examples underscore the potential of aptamers in cancer therapy and highlight the ongoing efforts in developing effective treatments.

Despite challenges and complexities, ongoing research aims to harness the therapeutic potential of aptamers to combat cancer effectively. Until very recently, SELEX was considered the golden standard method for the development of aptamers. In the past few years, however, we have demonstrated how it is possible to design aptamers specific for a given protein using computational tools^39^. For instance, we have developed *in silico de novo* RNA aptamers targeting TAR DNA-binding protein 43 (TDP-43), implicated in neurodegenerative diseases, demonstrating the strength and precision of an algorithm-based approach in the design of oligonucleotide sequences targeting a protein of interest^35,72^. Building on our previous studies in the computationally-driven development of aptamers, this work focuses on *in silico*-designed aptamers targeting RAD51. By leveraging these advanced computational tools, we can design drugs that target specific protein interactions with a level of precision that was previously unattainable.

The speed and accuracy of our computational approach allow for the rapid identification and optimization of aptamer sequences, effectively overcoming the limitations associated with conventional methods such as SELEX, which can be time-consuming and labor-intensive. This efficiency enables us to explore and exploit novel therapeutic targets like RAD51, opening up new avenues for drug development in areas where traditional small molecules have shown limitations in the context of formulation and delivery^73,74^. In the case of RAD51, our approach not only identified aptamers that specifically bind to the critical interaction sites but also provided a robust validation of their binding affinity and specificity through experimental techniques like TEM and FLIM.

In the TEM experiments, incremental addition of Apt1 to RAD51 determined the gradual disruption of the protein filaments. This interference of Apt1 with RAD51 quaternary conformation suggests a targeted mechanism by which this aptamer interacts with specific areas within the protein that are critical for its oligomerization. This is also in agreement with the AlphaFold 3 predictions of a RAD51 filament in the presence of an excess of Apt1. The predictions suggest that excess of Apt1 hinders the establishment of the typical inter-protomer interactions that usually lead to the formation of a stable filament, thus less likely resulting in a long fibril. Indeed, in the presence of many Apt1 copies, the thermodynamic equilibrium of the RAD51 oligomerization would probably shift towards the formation of short RAD51 oligomeric structures (e.g. trimers, hexamers) to the detriment of the helical filament. This is due to different interactions between alternative regions of the protein taking place or promoted by the DNA aptamer.

Since RAD51 plays a crucial role in HR, a process essential for the accurate repair of DNA double-strand breaks, the absence of RAD51 filaments could impair the protein’s ability to initiate and propagate the repair of damaged DNA, leading to an accumulation of unrepaired DNA damage in cancer cells. Such damage could potentially enhance the cytotoxic effects of DNA-damaging agents used in cancer therapy, making cancer cells more susceptible to treatment and less likely to recover and proliferate.

As shown by FLIM experiments, the displacement of Apt1 by the addition of BRC4 suggests that they either bind the same region of RAD51 or that the interaction of BRC4 with the protein determines a conformational rearrangement incompatible with the concurrent binding of the aptamer. These results demonstrate the power of our computational methods to create effective highly targeted aptamers that can selectively disrupt crucial protein-protein interactions, offering new possibilities for cancer treatment.

The effect of aptamers targeting RAD51 was investigated also in cellular contexts. In particular, we examined how Apt1 affects cellular DNA damage response and repair, focusing on HR through IF microscopy. We used γH2AX nuclear *foci* as markers for DSBs, crucial indicators of DNA damage and repair dynamics. Cells treated with Apt1 showed a significant increase in DSBs compared to untreated and negative control conditions, suggesting that tailored-designed aptamers against RAD51 may disrupt the physiological repair processes, potentially targeting and inhibiting the HR pathway, vital for accurate DSB repair. Moreover, the ability of Apt1 to inhibit the formation of RAD51-positive *foci* under stress-induced conditions indicate their potential to inhibit HR-mediated DNA repair pathways, further substantiates the role of this aptamer as potent disruptor of the RAD51-BRCA2 interaction. IF microscopy also showed that Apt1 reduced RAD51’s nuclear transport, inhibiting its recruitment to DNA damage sites. This finding suggests our aptamers’ potential to impair cancer cells’ reliance on HR for survival, making them a viable tool to be combined with other molecules that interfere with different DNA repair pathways, in order to achieve SL.

The potential of SL in BxPC-3 pancreatic adenocarcinoma cells was investigated by administering a treatment combination of Apt1 and the PARP inhibitor olaparib. Olaparib specifically targets BRCA-mutated cancer cells, which are particularly vulnerable to HR inhibition, a key component of SL. The dual inhibition of HR and PARP by this combination disrupts critical DNA repair pathways, enhancing the cytotoxicity of the treatment and potentially circumventing the resistance often seen with monotherapy. In this regard, our data indicate that Apt1 is able to inhibit RAD51-BRCA2 interaction, thus functionally mimicking the pathological phenotype of BRCA2-mutated cancer cells and, hence, sensitise their olaparib-mediated cell death. This is a direct translation of our previously shown paradigm of “fully chemically-induced synthetic lethality” from a small molecule-based to an aptamer-based strategy, paving the way for the development of novel pharmacological approaches.

Although aptamers exhibit high specificity and low immunogenicity, their transition from laboratory to clinical application is full of challenges, such as their potentially limited stability in the intricate cellular environment and *in vivo*. Our future efforts will be directed towards enhancing the delivery mechanisms of aptamers to ensure optimal bioavailability and therapeutic efficacy in clinical settings. Future investigations will also focus on elucidating the *in vivo* safety profiles of these aptamers, paving the way for their transition towards effective cancer therapies. Lastly, the exploration of combinatorial strategies involving aptamers and other cancer therapies will be further evaluated, since it could unlock synergistic effects with enhanced therapeutic outcomes, potentially transforming the field of cancer treatment across diverse cancer types.

In conclusion, this research provides groundbreaking insights into the design and functional characterization of aptamers targeting RAD51, shedding light on their potential as transformative therapeutic agents in cancer treatment by exploiting vulnerabilities in DNA repair mechanisms. Moreover, this novel study advances the development of nucleic acid-based drugs highlighting aptamers as exceptionally promising agents to amplify the efficacy of existing cancer treatments by interfering with DNA repair mechanisms, in the context of SL. By making cancer cells more vulnerable, this strategy would enable a reduced dosage of chemotherapy drugs while achieving the same therapeutic effect, significantly lowering the side effects associated with chemotherapy. Reducing these side effects is crucial for cancer patients, as it can substantially improve their quality of life during treatment. The path forward offers the potential to reveal innovative therapeutic approaches, possibly reshaping the field of cancer treatment and providing new potential outcomes to patients fighting against cancer.

## Material and Methods

### COMPUTATIONAL DESIGN OF APTAMERS

As a starting point for the design of the aptamer we retrieved ChIP-seq peaks for RAD51 available in the ENCODE project under the MCF-7 cancer cell line (https://www.encodeproject.org/experiments/ENCSR442VBJ/). Both Irreproducible Discovery Rate (IDR _thresholded peaks (foreground set) and background peaks used for IDR estimations were utilized. The foreground peaks were filtered for log10(q-value) ≥ 3, while the background peaks were filtered for log10(q-value) ≤ 0.4. From each set, nucleotide fragments of 100 nt and centered around the peak point-source were extracted, resulting in foreground and background sets of 3,157 and 67,263 nucleotide fragments, respectively. These two sets were used as input into the DREME algorithm from MEME Suite software (http://meme-suite.org/doc/dreme.html) to identify motifs relatively enriched in the foreground set compared to the background. DREME was configured with a minimum motif length of 4 and a maximum of 15, detecting enriched motifs corresponding to 114 RNA short sequences that were used to create aptamer sequences with length of approximately 16nt.

For the calculations, two RAD51 regions were defined (RAD51_Region 1, RAD51_Region 2) as protein fragments with length of ca. 70 amino acids and centered around the interaction pockets between RAD51 and a peptide derived from BRCA2 called BRC4^75,76^. The interaction sites are well defined and the BRC4 structure is well known^44,77,78^. Region 1 spans from Gly151 to Ile220, and region 2 spans from Asn196 to Ile265. A third RAD51 region of 70 amino acids, spanning from Glu29 to Glu98, was defined as Region 3, centered around the HhH domain (Thr48 to Glu77):

- RAD51_Region 1 (**Supplementary Fig. 1A**): GGGEGKAMYIDTEGTFRPERLLAVAERYGLSGSDVLDNVAYARAFNTDHQTQLLYQASAMMVESRYALLI
- RAD51_Region 2 (**Supplementary Fig. 1B**): NTDHQTQLLYQASAMMVESRYALLIVDSATALYRTDYSGRGELSARQMHLARFLRMLLRLADEFGVAVVI
- RAD51_Region 3 (**Supplementary Fig. 1C**): EQCGINANDVKKLEEAGFHTVEAVAYAPKKELINIKGISEAKADKILAEAAKLVPMGFTTATEFHQRRSE

As a control, we used fragments derived from actin B (Uniprot <u>P60709</u>), a highly expressed cytoskeletal protein in most cell lines. Using a sliding window approach with a 35 amino acids step and a window of 70 amino acids length, a control set of 9 fragments was generated. Then, *cat*RAPID algorithm ^38,39^ was employed to estimate the interaction propensities of the aptamer sequences against Region 1 and Region 2 compared to the control actin B protein fragments (*cat*RAPID score). For each aptamer, mean and standard deviation (SD) of interaction scores against the actin B control protein fragments were estimated. Aptamers were firstly filtered for *cat*RAPID scores exceeding the minimum threshold that is defined as half a standard deviation above the background mean (threshold=mean_background+0.5*standard_deviation_background; catRAPID_pocket>= threshold). Finally, the filtered aptamers showing a higher *cat*RAPID score against either Region 1 or Region 2 compared to Region 3, were considered as potential aptamers with specificity to RAD51 regions.

### COMPUTATIONAL PREDICTION OF APTAMER-PROTEIN COMPLEXES

The AlphaFold 3 deep-learning framework developed by Google DeepMind and Isomorphic Labs (available as a web server at https://alphafoldserver.com/about) was used to obtain a first *in silico* validation of the aptamers specificity for RAD51 pockets. This recently updated version of the AlphaFold AI model was chosen since it can generate structure predictions of hetero-complexes starting from one or multiple copies of different types of molecules, including proteins, peptides and nucleic acids, and demonstrated a much higher accuracy for protein-nucleic acid interactions over nucleic acid-specific predictors^45^.

By means of the AlphaFold Server, several combination of different molecules and different copy number were tested: RAD51 by itself (from Gly21 to Asp339) in 1, 6 and 7 copies; 1 or 6 RAD51 copies with 1, 6, 12 copies of aptamer (*i.e.* Apt1 and rcApt). For the sake of completeness, the structural prediction of RAD51 oligomeric filament was also run in the presence of both BRC4 (1, 3 or 6 copies) and Apt1 (**Supplementary Information**).

Each job (producing five different predicted models) was repeated five times on the platform using a different randomly generated seed, for each chosen condition.

Careful visual investigation of the models was performed to verify the quality of the predictions (*e.g.* absence of steric clashes, loop reconstruction). Given such inspections, the models presented in the **Results** (as well as in the **Supplementary Information**) represent the best prediction in terms of predicted template modeling score and interface predicted template modeling score, two of the typical AlphaFold 3 metrics.

For image rendering and contact analysis (reported in **Supplementary Table 2**), the calculations were run taking advantage of VMD 1.9.4 tools ^79^. Considering the absence of the hydrogens in the models, the threshold distance between the heavy atoms of the protein and the aptamer nucleotides was set at 4 Å in order to include all the putative electrostatic interactions.

### APTAMERS CHEMICAL CHARACTERISTICS

The aptamers here employed were purchased from Eurofins and ATDbio and display lengths varying between 12 to 15 bases. The aptamer called “rcApt” is the reverse complementary sequence of Apt1 and acts as a negative control in these studies. For the BLI experiments, biotin was attached to the 5’ end of all sequences, facilitating molecule binding and detection. For the in-cell assays, locked bases were introduced at both 5’ and 3’ ends of the aptamers Apt1 and rcApt, to increase their stability. In addition, the fluorescent dye Texas Red was conjugated to the 5’ end, providing a visual marker also for FLIM experiments. Prior to each experiment, whether conducted *in vitro* or in cell, the aptamers were subjected to an incubation period of one hour at 30°C to facilitate their correct folding.

### PROTEIN PRODUCTION AND PURIFICATION

The full-length human RAD51 protein modified with a 6×His tag at its N-terminus was produced in *E. coli* Rosetta2(DE3) pLysS cell strain using a pET15b plasmid, containing the gene for ampicillin resistance. The plasmid with the cDNA for 6xHis tag-RAD1 protein was inserted into the Rosetta2(DE3)pLysS by transforming the cells by means of thermal shock treatment, according to literature ^44^. A starter culture was prepared using a single colony of transformed Rosetta2(DE3)pLysS cells, incubated overnight at 37 °C in Luria-Bertani (LB) medium supplemented with 100 µg/mg ampicillin. The starter culture was then scaled up in appropriate volume by diluting it in 100-fold volume of fresh LB, always supplemented with 100 µg/ml ampicillin. During growth, flasks were shaken at 200 rpm and incubated at 37°C until the cell reached an optical density (OD_600_) of 0.6 - 0.8. At this point, Isopropyl ß-D-1-thiogalactopyranoside (IPTG) was added to the culture at a final concentration of 0.5 mM to induce heterologous protein production, and the cells were left to grow overnight at 37 °C, while shaking. Cells were collected via centrifugation at 3000 g at 4°C for 20 minutes. The pellet was resuspended in a suitable volume (10 mL/gram of wet cell mass) of lysis buffer containing 20 mM Tris-HCl at pH 8.00, 500 mM NaCl, 10 mM imidazole, 5 mM 1,4-Dithiothreitol (DTT) and 10% glycerol by volume, supplemented with a protease inhibitor mix (Protease Inhibitor Cocktail Tablets, EDTA-Free, Sigmafast) and DNAse I (from Merck). Cell lysis was achieved through sonication on ice (14 cycles of 30 seconds each, and amplitude set at 40%) using a Bandelin Sonoplus HD2070 immersion sonicator. The lysed cell mixture was then centrifuged at 18,000 g at 4°C for 1 hour. Subsequently, the protein extract (supernatant) was introduced into a column packed with nickel-loaded resin, previously equilibrated with buffer A1, which corresponds to lysis buffer without protease inhibitor and DNase I. After the flow-through was discarded, proteins without the His-tag were washed away with a wash buffer (20 mM Tris-HCl, pH 8.00, 500 mM NaCl, 25 mM imidazole, 10% (v/v) glycerol). The target protein was eluted with the same buffer containing 75 mM imidazole. The eluted protein solution was dialysed overnight at 4°C in buffer A2 (50 mM Tris-HCl, pH 8.00, 200 mM KCl, 0.25 mM EDTA, 2 mM DTT, 10% (v/v) glycerol) using a dialysis membrane (CarlRoth Cat. N 1991.1) with a cut-off of 10 kDa and then processed through an anion exchange column (ResQ, GE Healthcare) pre-equilibrated with buffer A2. Elution was carried out using a gradient of buffer B2 (50 mM Tris-HCl, pH 8.00, 1 M KCl, 0.25 mM EDTA, 2 mM DTT, 10% (v/v) glycerol), reaching 100% B2 in 10 column volumes. High purity RAD51-containing fractions were pooled and dialysed against storage buffer (20 mM Hepes, pH 8.00, 250 mM KCl, 0.1 mM EDTA, 2 mM DTT, 10% (v/v) glycerol) overnight at 4°. Protein yield was verified using the optical absorption at 280 nm (extinction coefficient 14,900 M^−1^cm^−1^) of the final sample and was determined to be 200 ng/ml on average. Protein aliquots were flash-frozen and stored at –80 °C.

### BIOLAYER INTERFEROMETRY

BLI was employed to determine binding constants (K_d_s) of the aptamers with RAD51. Experiments were conducted using an Octet Red instrument (ForteBio, Inc., Menlo Park, CA) set at 25 °C. All steps of the assay were executed in binding buffer, a solution of 50 mM Tris HCl buffer at pH 8, containing 200 mM KCl, 0.25 mM EDTA, 1 mM DTT, 10% glycerol and 0.01% Tween-20. Biosensors coated with streptavidin were selected to enable the loading of biotinylated DNA aptamers at a concentration of 2 μg/ml. The protocol to generate the binding curves between the aptamers and RAD51 was set as follows: 180-second baseline; 300-second aptamer loading step on the sensors; 120-second washing step; 600-second association step with increasing protein concentrations (ranging from 20 nM to 20 μM, according to the binding strength); 600-second dissociation step. The experiments were performed with constant shaking at 250 rpm. K_d_ values were derived by fitting the response intensity (shift in wavelength upon binding) against the protein concentration at a steady state. The experiments were performed in triplicates.

### NEGATIVE STAINING TRANSMISSION ELECTRON MICROSCOPY

Recombinant RAD51 protein (2.5 μM) was incubated at 4°C for 1 hour, either by itself or in the presence of increasing concentrations of Apt1. After incubation, each sample (5 µL) was adsorbed onto pure carbon film 300-mesh copper grids (Electron Microscopy Sciences, Hatfield, PA, USA). Following several washes in the glycerol-free storage buffer (20 mM Hepes, pH 8.00, 250 mM KCl, 0.1 mM EDTA, 2 mM DTT), each sample was negatively stained using 1% uranyl acetate in Milli-Q water. The samples were observed using a JEM-1011 (JEOL) transmission electron microscope (TEM) equipped with a thermionic source (W filament) and a maximum acceleration voltage of 100 kV. The microscope was fitted with a Gatan Orius SC1000 series CCD camera (4008 × 2672 active pixels), fiber optically coupled to a high-resolution phosphor scintillator. The average diameter of the ring-like structures, identified in the RAD51:Apt1=1:6 condition, was calculated using the tool analyze and measure of the image analysis software ImageJ.

### FLUORESCENCE-LIFETIME IMAGING MICROSCOPY

FLIM images have been acquired using a Digital Frequency Domain (DFD) FLIMbox (ISS inc., Champaign, IL) coupled to a A1RMP multiphoton microscope (Nikon, Japan), calibrated with a 10 µM solution of Coumarin 153 in methanol (τ= 4,3 ns). Samples were 2-photon excited focusing a Chameleon UltraII Ti:Sapphire laser (Coherent inc.,Saxonburg, PA) tuned at 850 nm through a 60x oil objective (NA=1.45). Fluorescence emitted photons were filtered using a 525/50 nm BP filter for Coumarin 153 and a 605/70 nm for Texas Red labeled aptamers. The analyses were performed with ISS VistaVision multi-image phasor analysis software, where sine (s) and cosine (g) coordinates are defined as

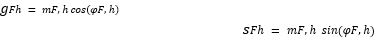

Where *m* is the modulation and *φ* the phase acquired of the *h*-th pixel in the FLIM image and *F* the laser repetition rate.

### CELL CULTURE HANDLING

The human pancreatic ductal adenocarcinoma BxPC-3 cell line was obtained from American Type Culture Collection (ATCC; Manassas, VA, USA). BxPC-3 cells were grown in RPMI 1640 medium (Thermofisher, 11875093), supplemented with 2 mM L-glutamine, 100 U/mL penicillin, 100 U/mL streptomycin, and 10% fetal bovine serum (FBS). Cells were maintained at 37°C in a humidified 5% CO_2_ atmosphere and split when at confluence. For microscopy analysis, cells were seeded at a concentration of 150000 cells/ml onto 24-well plates with coverslips. After a day, complete medium was replaced with FBS-free medium for 18h-24h, to synchronize the cells to the same cell cycle phase. Subsequently, fresh complete medium was introduced, and, after a 2-hour recovery period, cells were treated with Lipofectamine 2000 (Thermofisher, 11668019) with or without 0.5 μg/ml aptamers, following manufacturer instructions. Following transfection, cells were allowed to recover overnight before further treatments. All culture media and supplements, unless otherwise specified, were obtained from Sigma Aldrich (St. Louis, MO, USA). Olaparib (PubChem CID: 23725625, #S1060) and *cis*-diamminedichloroplatinum(II) (cisplatin) (PubChem CID: 2767, #S1166) were purchased from Selleck Chemicals (Houston, TX, USA). Olaparib was dissolved in DMSO at the final concentration of 10 mM. Cisplatin was dissolved in 1X PBS at the final concentration of 5 mM. Stock aliquots were stored at -20° C and diluted in complete medium prior to each experiment.

### IMMUNOFLUORESCENCE SAMPLE PREPARATION FOR RAD51 BEHAVIOR INVESTIGATION

The day after transfection, half of the sample cells were treated with 50 µM cisplatin for 2 hours to induce DNA damage, while the other half was left untreated as a control. Following cisplatin treatment, cells were rinsed two times with phosphate-buffered saline (PBS) and then fixed for 10 minutes at room temperature using a 4% paraformaldehyde solution prepared in PBS. Subsequent to three PBS washes, cells underwent permeabilization with 0.1% Triton-X 100 in PBS for a duration of 5 minutes, followed by three more PBS washes.

Cells were treated with 5% BSA prepared in PBS for 30 minutes as a blocking step. Following this, cells were exposed to a primary antibody to detect the RAD51 protein using rabbit anti-RAD51 antibody (Bio Academia, 70-001) prepared in 5% bovine serum albumin (BSA) in PBS at a 1:1000 dilution, and incubated for 1 hour at room temperature. Subsequently, a secondary anti-rabbit antibody conjugated to Alexa Fluor 488 (Thermofisher, A-11008) was added at a 1:1000 dilution and incubated for an additional hour at room temperature.

After three washes with PBS, cells were treated for 5 minutes with a 0.5 μg/ml solution of 4′,6-diamidino-2-phenylindole (DAPI) prepared in PBS. Following three additional rinses in PBS, the coverslips were positioned face-down on glass slides using ProLongTM Diamond Antifade Mountant (Thermofisher, P36965).

### IMMUNOFLUORESCENCE SAMPLE PREPARATION FOR DNA DAMAGE QUANTIFICATION

To evaluate the effect of aptamers on physiological DSBs, cells were analyzed three days after transfection by assessing the ones positive for γH2AX *foci*. To determine the aptamer impact on SL, olaparib was added at a concentration of 10 µM one day post aptamer transfection. The formation γH2AX *foci* was analyzed three days after transfection. In both cases cells were rinsed two times with PBS and then fixed for 10 minutes at room temperature using a 4% paraformaldehyde solution prepared in PBS. Subsequent to three PBS washes, cells underwent permeabilization with 0.1% Triton-X 100 in PBS for a duration of 5 minutes, followed by three more PBS washes. Cells were treated with 5% BSA prepared in PBS for 30 minutes as a blocking step. Following this, a primary antibody against phospho-histone H2AX (Ser139), clone JBW301 (Merck/Sigma Aldrich, 05-636), was employed at a 1:1000 dilution in 5% BSA/PBS and incubated for 1 hour at room temperature. Subsequently, cells were treated with a secondary anti-mouse antibody conjugated to Alexa Fluor 647 (Thermofisher, A-21236) at a 1:1000 dilution and incubated for an additional hour at room temperature.

After three washes with PBS, cells were treated for 5 min with a 0.5 μg/ml solution of 4′,6-diamidino-2-phenylindole (DAPI) prepared in PBS. Following three additional rinses in PBS, the coverslips were positioned face-down on glass slides using ProLongTM Diamond Antifade Mountant (Thermofisher, P36965).

### IMMUNOFLUORESCENCE IMAGES ACQUISITION

Slides containing the fixed cells were examined using a Nikon’s A1R confocal microscope, using the objective Plan Apo TIRF 60x/1.49 Oil DIC H N2 along with 4-channel detectors and by means of the Nikon software NIS-Elements Advanced Research version 5.30.02 (64-bit). Fluorescence patterns were determined by sketching a line over selected cells and assessing the fluorescence intensity of the fluorophores associated with the molecule to be detected (DAPI, RAD51, γH2AX, and TexasRed) on a pixel-by-pixel basis. Images were acquired with a pixel size of 104 nm by maintaining the fluorescence settings constant, to remove any potential bias. At least 8 images per condition were examined. A minimum of 600 cells per sample were used to compute fluorescence intensities and *foci* counts.

### IMMUNOFLUORESCENCE IMAGES ANALYSIS

Our in-house macro was run on the software ImageJ to perform the RAD51 *foci* count with and without cisplatin and in the absence/presence of fluorophore-tagged locked nucleic acid (LNA) Apt1 or rcApt, to assess the impact of aptamers on DSB repair mechanism. To measure the intensity of nuclear RAD51 fluorescence, the nuclear region was identified using the nuclear stain DAPI. To segment the nuclei, a Gaussian filtering was applied to the images to smooth them and reduce noise. The image threshold was adjusted (Otsu method) to differentiate between the signal of interest and background. Finally, a watershed algorithm was applied to the binary image to separate *nuclei* that might be touching each other. Particle analysis parameters were set as follows: size range 1 to infinity; circularity between 0.5 and 1. Once identified in the nuclear region and added to the region of interest (ROI) manager, nuclear fluorescence was quantified in the green channel, corresponding to RAD51. The fluorescence data were normalized by nuclear area. To estimate the fluorescence intensity ratio between the cytoplasmic and nuclear RAD51 signal, a macro was designed for ImageJ with the following operational pipeline. The programme was configured to measure the area and the mode of fluorescence intensity of ROIs representing nuclear and cytoplasmic areas in the images. A Gaussian blur with Sigma 1 was applied to the DAPI channel, aiding in the segmentation of the nuclear region with higher confidence. The image was segmented using the Otsu criterion, an automatic and robust threshold parameter. Holes in the binary images were filled using the ImageJ algorithm, and these were used to create a region of interest.

The second channel -the signal from RAD51- was processed in a similar manner. A Gaussian blur with sigma 0.5 was applied, and the image was segmented using the MinError function; this allowed for the obtaining of a binary image of the entire cell due to the RAD51 distribution. The binary image of the nucleus was subtracted from that of the whole cells to obtain the cytoplasmic area image. Finally, the mode intensity and area of the two regions of interest, the cytoplasmic and nuclear, were measured. The ratio between the nuclear and cytoplasmic signal was used to discriminate between control and treated cells as previously described. The fluorescence values of nuclear RAD51 are calculated by dividing the nuclear mean fluorescence by the nuclear area and are reported as percentage. Mean/area of cells treated with cisplatin only is considered to be 100%.

To assess the increased presence of γH2AX *foci* within nuclei due to DNA damage, a macro was developed for ImageJ, establishing a comprehensive operational protocol for automated image analysis across a specified directory. Image parameters, such as the directory path, file format, and channels designated for segmentation and analysis, were configured at the outset. A reference image, specifically one of cells treated with cisplatin, was selected to adjust threshold parameters essential for segmenting the signal of interest. Subsequent to this initial setup, a Gaussian blur with a sigma value of 1 was applied to the DAPI channel to enhance the segmentation accuracy of the nuclear regions. The segmentation was executed employing the Otsu criterion, an automated and robust thresholding method. Holes within the binary images were filled using ImageJ’s algorithm, facilitating the delineation of individual nuclei as regions of interest (ROIs) through the Analyze Particles function. Parameters were set to include objects with a size of at least 5000 pixels, a circularity range of 0.5-1.00, and to exclude objects at the image periphery.For the analysis of the γH2AX signal, a Gaussian blur with a sigma of 0.5 was applied, followed by segmentation using threshold values determined from the reference image. This process generated a binary image of the γH2AX *foci*. Each nuclear ROI containing *foci* was quantified by calculating the mean pixel value within the binary γH2AX image; a mean exceeding one was indicative of a positive ROI. The proportion of positive *nuclei* to the total number of *nuclei* served to distinguish between control and treated cells as delineated in the study.

### CELL VIABILITY ASSESSMENT

Cell viability was assessed via MTT [3-(4, 5-dimethylthiazol-2-yl)-2,5-diphenyltetrazolium bromide] assay as described previously^80^. BxPC-3 cells (150,000 cells/mL) were seeded at a volume of 500 μL per well in 24-well plates and allowed to adhere overnight. After 24 hours, cell transfection with 0.5 μg/mL aptamers and Lipofectamine® 2000 was carried out as previously described, following the manufacturer’s instructions. After a 4-hour incubation, the transfection medium was replaced with fresh complete medium, and cells were incubated at 37°C overnight. The following day, cells were treated with 10 μM olaparib to assess synthetic lethality upon inhibition of the RAD51-BRCA2 interaction. After a 72-hour treatment with olaparib, a sterile solution of 1 mg/mL MTT in 1X PBS was added to each well at a final concentration of 0.1 mg/mL. The plates were incubated at 37°C for 4 hours, and the formazan crystals were solubilised overnight by adding a 1:1 volume of 10% SDS/0.01M HCl solution to each well. Absorbance was measured at 570 nm and 690 nm wavelengths.

## Acknowledgements

The authors would like to thank the other members of Tartaglia and Cavalli’s groups. The research leading to this work was supported by the ERC ASTRA_855923 (G.G.T.), EIC Pathfinder IVBM4PAP_101098989 (G.G.T.) and PNRR grant from National Centre for Gene Therapy and Drugs based on RNA Technology (CN00000041 EPNRRCN3 (G.G.T.). F.D.P. acknowledges the support granted by the European Union - Next Generation EU, Mission 4 Component 1 CUP D53D23016360001, PRIN-PNRR, Grant n. P2022CLXMK, for his current position. L.B. is supported by a Marie Skłodowska-Curie post-doctoral fellowship (UNDERPIN_101063903). The authors extend their gratitude to Giuseppe Vicidomini and Eleonora Perego for their help with the microscopy experiments. The authors acknowledge also the fundamental support of the Nikon Imaging Centre and the Electron Microscopy facility at Fondazione Istituto Italiano di Tecnologia.

## Author contributions

GM performed the experiments, participated in their design and wrote the manuscript. EZ performed and supervised the experiments and wrote the manuscript. AA generated the aptamers with GGT. FDP carried out the structural predictions and analysis of RAD51 complexed with aptamers and wrote the manuscript. MM supported, supervised the cell experiments and carried out the SL experiment. MG and LB provided assistance with cellular experiments. JR contributed to *in vitro* experimental work. GV was involved in producing some of the recombinant protein. MO developed the macro used for in-cell analysis. MS worked on the FLIM experiment. RM worked on the TEM experiment. SG designed and supervised the experiments. AC formulated the initial hypothesis and designed the experiments. GGT formulated the initial hypothesis, supervised the work and wrote the manuscript.

## Additional information

Additional information can be found in the Supplementary Materials.

