## Supplementary Information for "Computationally-designed aptamers targeting RAD51-BRCA2 interaction inhibit RAD51 nuclear recruitment"

### Tables

| ROI | Measurements |
|---|---|
| 1 | 13,07 |
| 2 | 11,36 |
| 3 | 13,07 |
| 4 | 10,80 |
| 5 | 15,34 |
| 6 | 13,64 |
| 7 | 14,77 |
| 8 | 14,20 |
| 9 | 13,64 |
| 10 | 13,07 |
| 11 | 14,20 |
| 12 | 13,64 |
| 13 | 11,93 |
| 14 | 12,50 |
| 15 | 11,93 |
| 16 | 17,05 |
| 17 | 12,50 |
| 18 | 11,93 |
| 19 | 11,36 |
| 20 | 11,93 |
| 21 | 10,23 |
| 22 | 12,50 |

| | |
|---|---|
| 23 | 14,20 |
| 24 | 14,77 |
| 25 | 14,20 |
| 26 | 13,64 |
| 27 | 13,64 |
| 28 | 12,50 |
| 29 | 13,07 |
| 30 | 13,07 |
| 31 | 12,50 |
| 32 | 14,77 |
| 33 | 13,64 |
| 34 | 13,07 |
| 35 | 12,50 |
| 36 | 11,93 |
| 37 | 13,64 |
| 38 | 16,48 |
| 39 | 13,64 |
| 40 | 13,07 |
| 41 | 14,77 |
| 42 | 12,50 |
| 43 | 13,64 |
| 44 | 13,64 |
| 45 | 15,34 |
| 46 | 14,77 |
| 47 | 14,77 |
| 48 | 13,07 |
| 49 | 14,20 |
| 50 | 10,80 |

**Supplementary Table 1**. Table displaying the diameter measurements of annular-like structures formed by RAD51 protein when combined with a six-fold molar excess of Apt1.

| Filament:aptamer ratio | 1:1 model | | | | | | 1:6 model | | | | | | 1:12 model | | | | | |
|---|---|---|---|---|---|---|---|---|---|---|---|---|---|---|---|---|---|---|
| RAD51 aa | A | B | C | D | E | F | A | B | C | D | E | F | A | B | C | D | E | F |
| Lys111 | | | | | | | | | | | | | | | | | x | |
| Leu112 | | | | | | | | | | | | | | | | | x | |
| Glu128 | | | | | | | x | x | x | | x | x | | | | x | | |
| Phe129 | | | | | | | | | | | | | | | | x | x | |
| Arg130 | | | | | | | | x | x | | x | | | | | x | x | |
| Thr131 | | | | | | | | | | | | | | | | | x | |
| Lys205 | | | | | | | x | | | | | | | | | | | |
| Gln206 | | | | | | | x | | | | | | | | | | | |
| Ala209 | | | | | | | x | | | | | | | | | | | |
| Arg229 | x | x | x | x | | | x | x | x | | | | | | x | | | |
| Tyr232 | | | | | | | x | | | | | | | | | | | |
| Ser233 | | | | | | | x | x | x | | x | | | | | | | |
| Gly234 | | | | | | | x | x | x | | x | x | | x | | | | |
| Arg235 | x | x | x | x | | | x | x | x | x | x | x | x | x | x | | | |
| Gly236 | | | | | | | x | x | x | x | x | | | | x | | | |
| Leu238 | x | x | x | x | x | | x | x | x | x | x | x | x | x | | | | |
| Ser239 | x | x | x | x | x | | x | x | x | x | x | x | | x | x | | | x |
| Arg241 | x | x | x | x | x | | | | | | | | x | x | | | | |
| Gln242 | x | x | x | x | x | | | | | | | | x | x | | | | |
| Met243 | x | | x | | x | | | | | | | | | x | | | | x |
| Val269 | | | | | | | | | | | | | | | x | | | |
| Val270 | x | x | x | | | | | | | | | | | | x | | | |
| Ala271 | x | x | x | x | | | x | x | x | x | x | x | x | x | | x | | |
| Gln272 | | | | | | | | x | x | x | x | x | x | x | | x | | |
| Val273 | x | x | x | x | | | | x | x | x | x | | x | x | x | x | | x |
| Asp274 | x | x | x | x | | | | x | x | | x | x | | x | | x | x | x |
| Gly275 | | | | | | | | | | | | | | x | | x | | x |
| Ala276 | | | | | | | | | | | | | | | | x | | |
| Ala277 | | | | | | | | | | | | | | | | | | x |
| Met278 | x | x | x | | | | | | x | | | | | x | x | | | x |
| Phe279 | | | | | | | x | x | x | x | x | x | x | | x | x | | |
| Ala280 | | | | | | | x | x | x | x | | x | | | x | x | | |
| Ala281 | | | | | | | x | x | x | x | | x | | | x | | x | |
| Asp282 | | | | | | | x | | x | x | | | | | | | x | |
| Pro283 | | | | | | | x | | x | x | | | | x | | | | x |
| Lys284 | | | | | | | x | x | x | x | x | | | x | | x | | |
| Lys285 | x | | x | x | | | | | | | x | | | | x | | | |

| | | | | | | | | | | | | | | | | | | |
|---|---|---|---|---|---|---|---|---|---|---|---|---|---|---|---|---|---|---|
| Ile287 | x | x | x | | | | | | | | | | | | x | | | |
| Gly288 | x | x | x | x | x | | | | | | | | x | x | x | | | |
| Gly289 | x | x | x | x | x | | | | | | | | x | x | x | | | |
| Asn290 | x | x | x | x | x | | x | x | x | x | | x | x | | | | | |
| Ile291 | x | x | x | x | x | | | | | | | | x | x | | | | |
| Arg303 | | | | | | | | | | | | | | x | | | | |
| Lys304 | | | | | | | x | x | x | | x | | | x | | | | |
| Gly305 | | | | | | | x | | | x | | | | x | | | | |
| Arg306 | | | | | | | x | | x | x | | | x | x | | | | x |
| Gly307 | | | | | | | | | | | | | x | | | | | x |
| Glu308 | | | | | | | | | | | | | | | | | | x |
| Lys313 | | | | | | | | | x | | | | | | | | | |
| Tyr315 | | | | | | | | | x | x | | | | | | | | |
| Glu322 | | | | | | | | | | | | | | | | x | | |
| Asn330 | | | | | | | | | | | | | | | | | | x |
| Ala331 | | | | | | | | | | | | | | | | | | x |
| Ala337 | | | | | | | x | | | | | | | | | | | |
| Lys338 | | | | | | | x | | | | | | | | | | | |
| Asp339 | | | | | | | x | | | | | | | | | | | |

**Supplementary Table 2**. Summary of RAD51 amino acids, by chain, interacting (within 4 Å) with DNA nucleotides in the RAD51-Apt1 predicted models at different filament:aptamer ratio. The grey-shaded cells mark the residues pertaining also to RAD51 Region 2.

### FIGURES

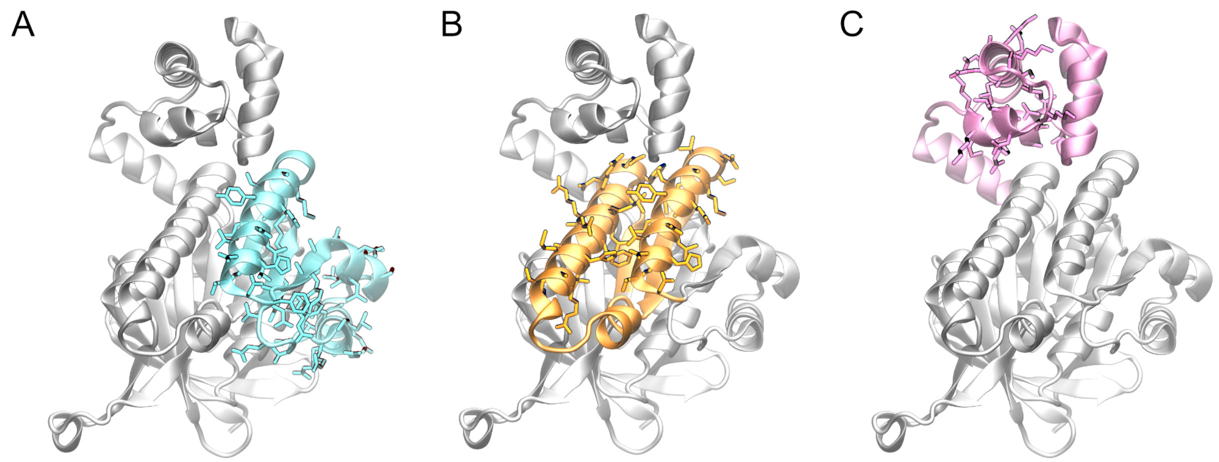


**Supplementary Figure 1. RAD51 structure and its regions of interest.** Cartoon representation of the single RAD51 protomer structure, similar to the incomplete X-ray structure of RAD51 deposited in the PDB with ID 1N0W[1]. The non-resolved elements – i.e. N-terminal domain (Gly21-Ser97), the L1 defined by Pellegrini et. al[1] connecting β-strand B4 to α-helix A5 (Thr230-Gly236), and L2 (Gln268-Ile292) between B5 and B6 – were reconstructed by means of Alphafold 3[2] prediction. **A**: Region 1 is highlighted in cyan, interacting with BRC4 motif FXXA, corresponding to the amino acids Ala157-Met210 on RAD51, represented in licorice; **B**: Region 2 is highlighted in orange, interacting with BRC4 motif LFDE, corresponding to the RAD51 residues Asn196-Ser214 and Ser239-Gly260, represented in licorice; **C**: Region 3, centered around the HhH domain (in licorice Thr48-Glu77), is highlighted in mauve.

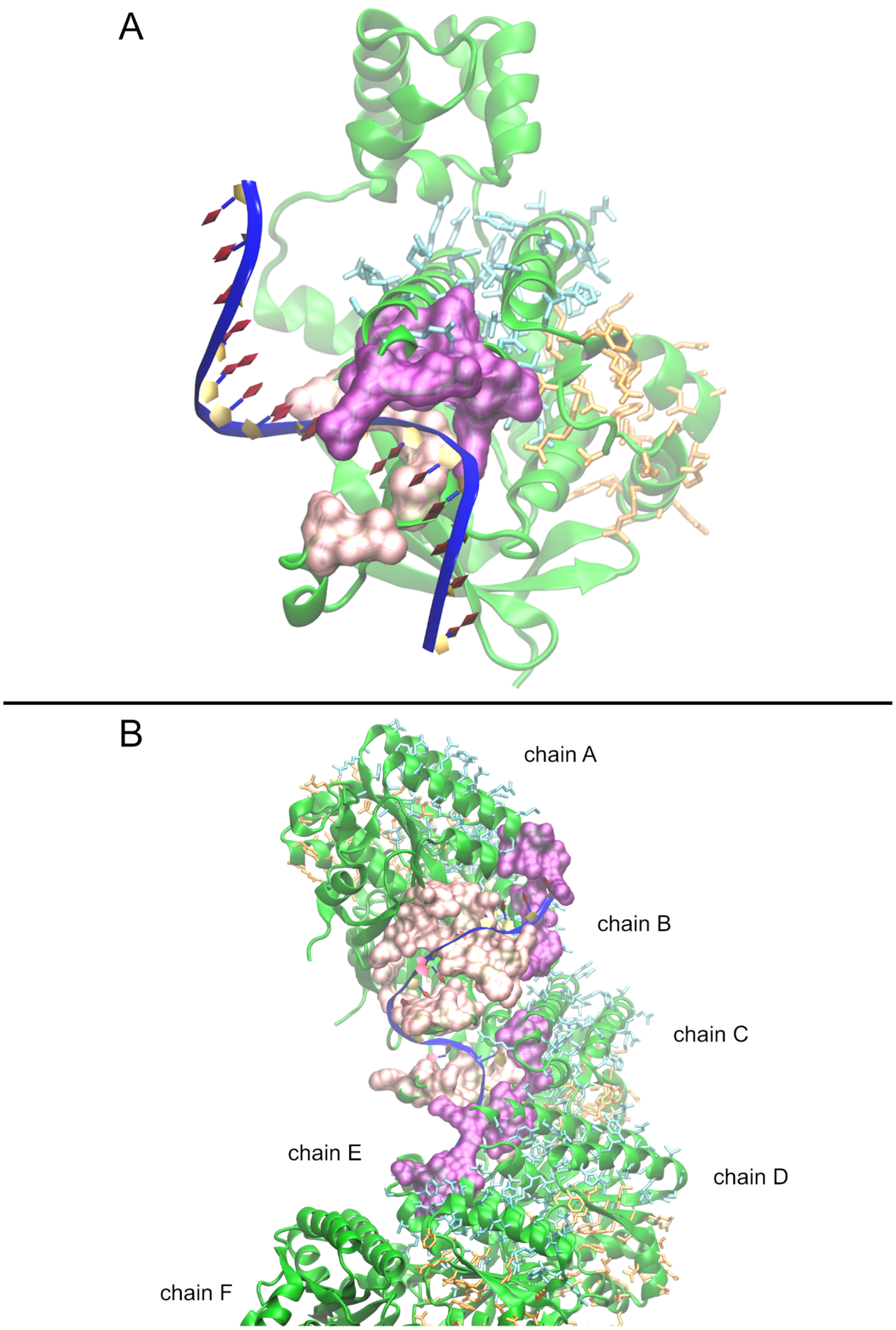


**Supplementary Figure 2. Focus on the RAD51/Apt1 interaction. A**: Single RAD51 protomer in complex with Apt1; **B**: RAD51 filament comprising 6 protomers (each N-terminal domain from residue 21 to 96 is omitted in the image but present in the prediction, for clarity purposes, to ensure that the residues pertaining to Region 1 and 2, shown in cyan and orange licorice, respectively, are not partially hidden) interacting with Apt1. This structure is analogous to that shown in **Fig. 4B**, with a 1:1 ratio, but +90° rotated). The aptamer is reported as a blue ribbon, with sugar rings in yellow and nucleobases in red. Each protein chain is shown in a green cartoon-based representation; chain F, not directly interacting with the DNA aptamer, is partially omitted to zoom in the surfaces shaping the binding groove. Residues in Region 1 and Region 2 are highlighted as licorice in orange and cyan, respectively. Violet surfaces indicate residues that belong to both Region 2 and the interacting amino acids with Apt1 nucleotides (within 4 Å), pink surfaces represent the residues in contact with the aptamer, but not included in Region 2. The combination of these two surfaces shapes the groove in which is lying the aptamer.

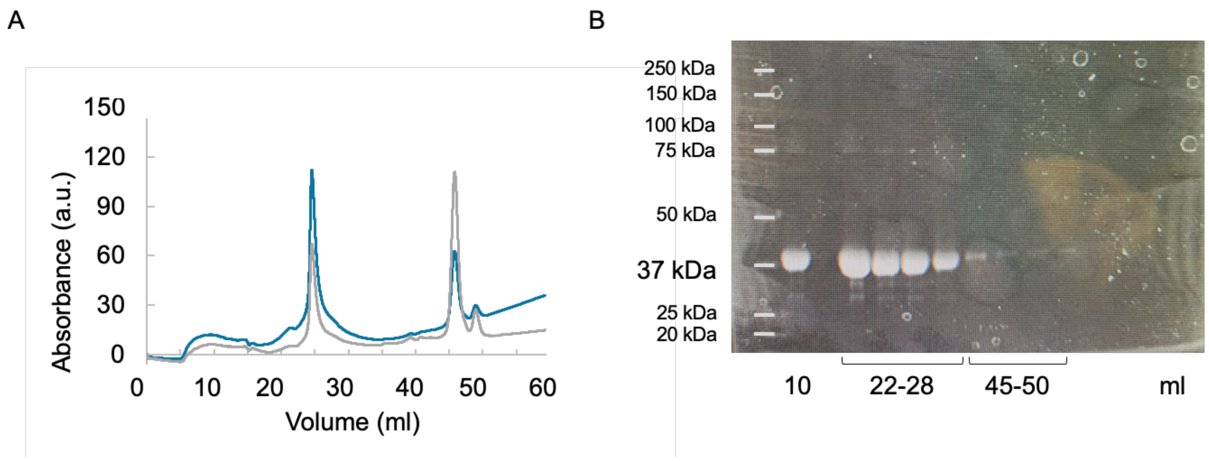


**Supplementary Figure 3: A** Chromatogram depicting the final step of RAD51 purification. Absorbance was recorded at 280 nm (shown in blue) and at 260 nm (shown in grey), with the protein eluting between volumes of 22-28 ml. **B**: SDS-PAGE illustrating the purity level obtained from the isolation of RAD51 protein following purification protocol.

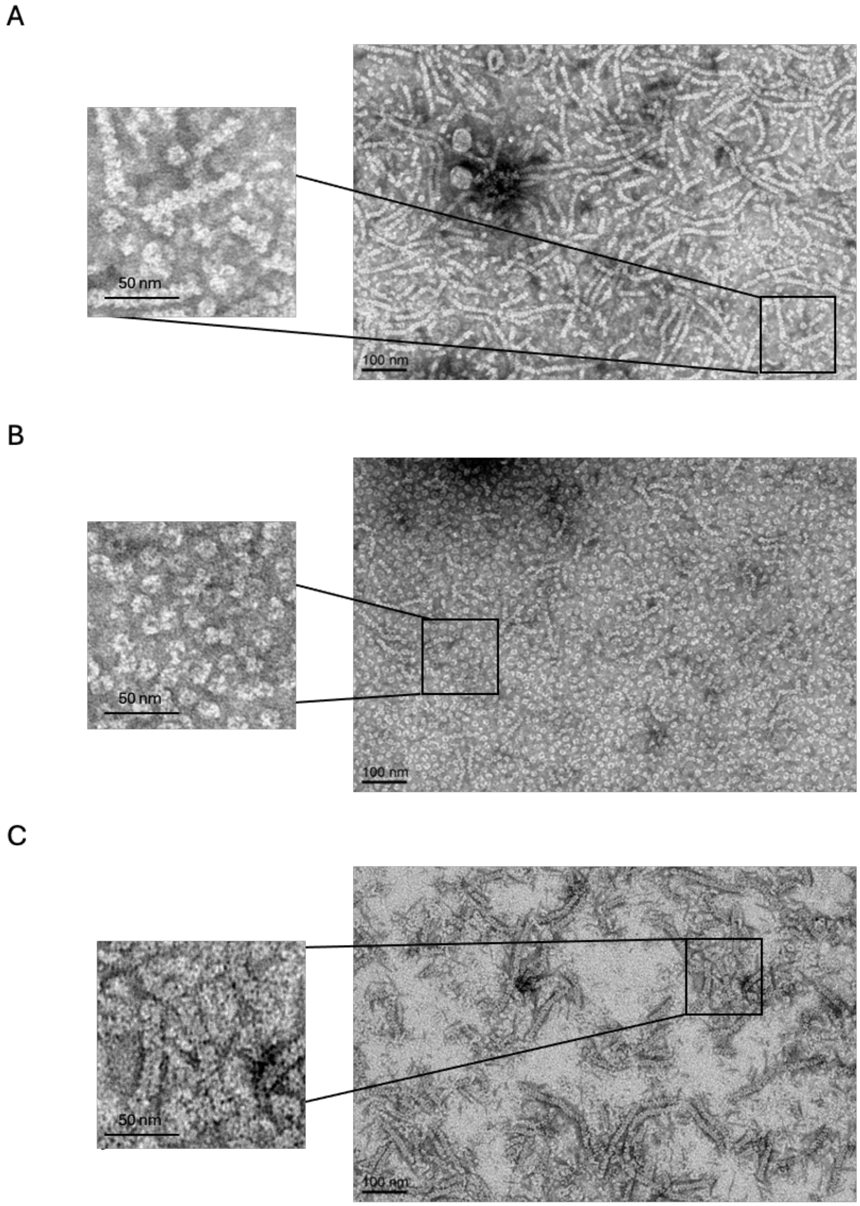


**Supplementary Figure 4: Negative staining TEM image of RAD51 in the presence of Apt1 and rcApt. A**: RAD51 incubated with Apt1 at a molar ratio of 1:3, showing the characteristic RAD51 fibril structure. This Apt1 concentration shows no alteration in the RAD51 fibril structures; **B**: RAD51 incubated with Apt1 at a molar ratio of 1:8, showing the peculiar RAD51 oligomers ring-like arrangement; **C**: RAD51 incubated with rcApt at a molar ratio of 1:12, showing the characteristic RAD51 fibril structure. The excess of negative control rcApt shows no alteration in the RAD51 fibril structures, confirming that the fibrils remain unchanged compared to RAD51 alone. This indicates that the negative control does not mimic the activity of Apt1, thereby validating Apt1's functionality.

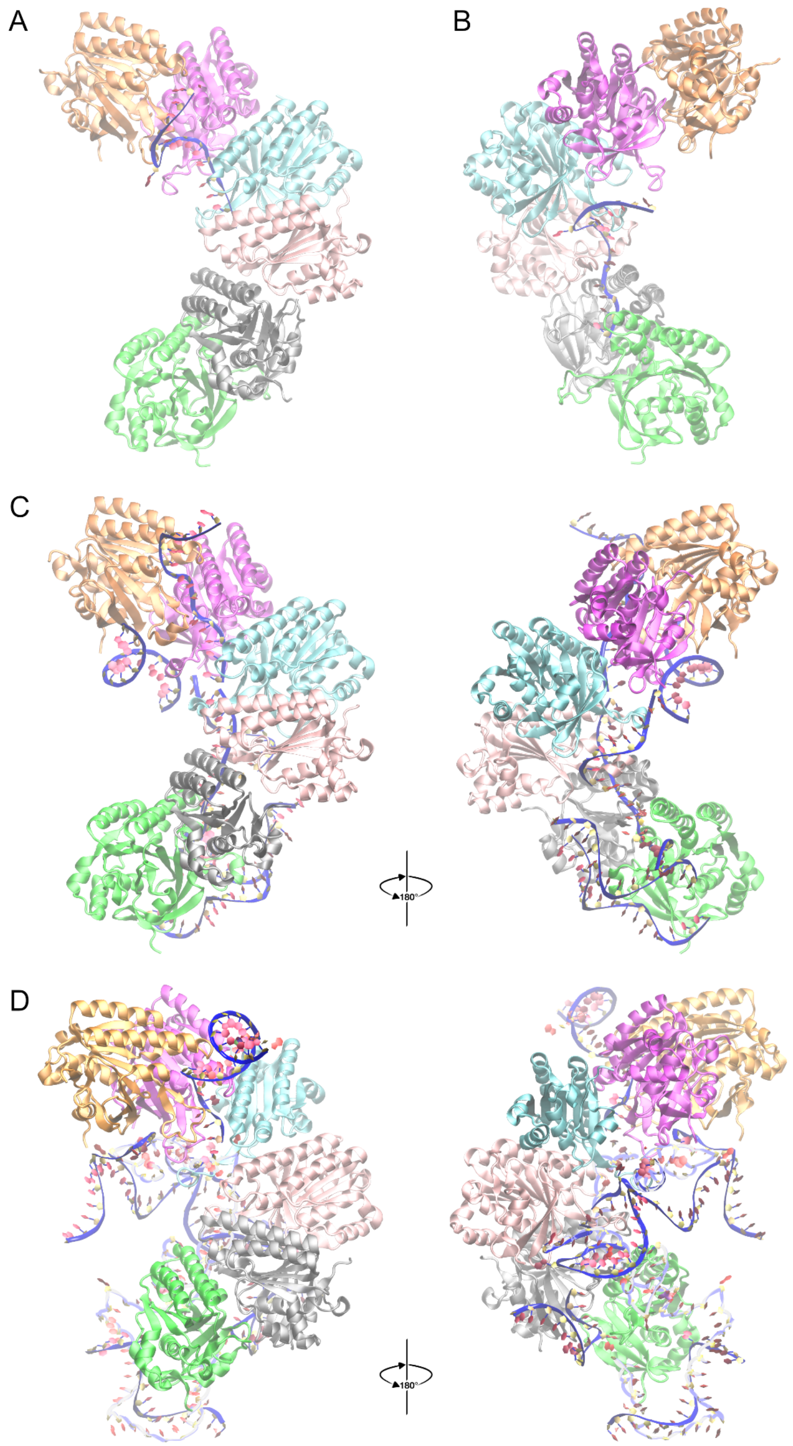


**Supplementary Figure 5. Structural prediction of RAD51 fibrils/oligomers in the presence of rcApt (control). A**: Single RAD51 filament (6 protomers), displaying the characteristic helical structure in complex with rcApt at a ratio of 1:1; **B**: Another possible model of RAD51 fibril in complex with rcApt at a 1:1 ratio, featuring the same fibril arrangement as in A, but with the aptamer in an alternative interaction; **C**: RAD51 fibril in complex with rcApt at a 1:6 ratio (together with its 180° rotated view). Unlike the RAD51/Apt1 complex at the same ratio (**Fig. 3C**), the fibril is still properly formed even in the presence of six rcApt copies: two of them are in the same binding pose as in panel A and B, instead the other four are non-specifically contacting the filament; **D**: RAD51 fibril in complex with rcApt at a 1:12 ratio (together with its 180° rotated view), even with double the number of rcApt copies, the fibril maintains maintains its typical helical shape with the DNA aptamers engaging in a non-recurrent, non-specific and non-conserved binding mode with the proteins. Thus, by means of *in silico* structural prediction, differently from Apt1, the rcApt does not interfere with the RAD51 filament formation. Indeed, 17/25 predictions were found in a *holo*-like conformation or binding rcApt in a bent conformation partially inserting one end in the ssDNA-bs (like panel A and B) or exactly in the RAD51 filament known ssDNA-bs, whereas 8/25 models were in a apo-like conformation not binding the rcApt at all or showing a week/partial non-specific binding. Each protein chain is shown in a different color using the cartoon-based representation (chain A in orange, B in magenta, C in cyan, D in pink, E in silver and F in green). Each aptamer copy is reported as a blue ribbon, with the sugar rings in yellow and the nucleobases in red.

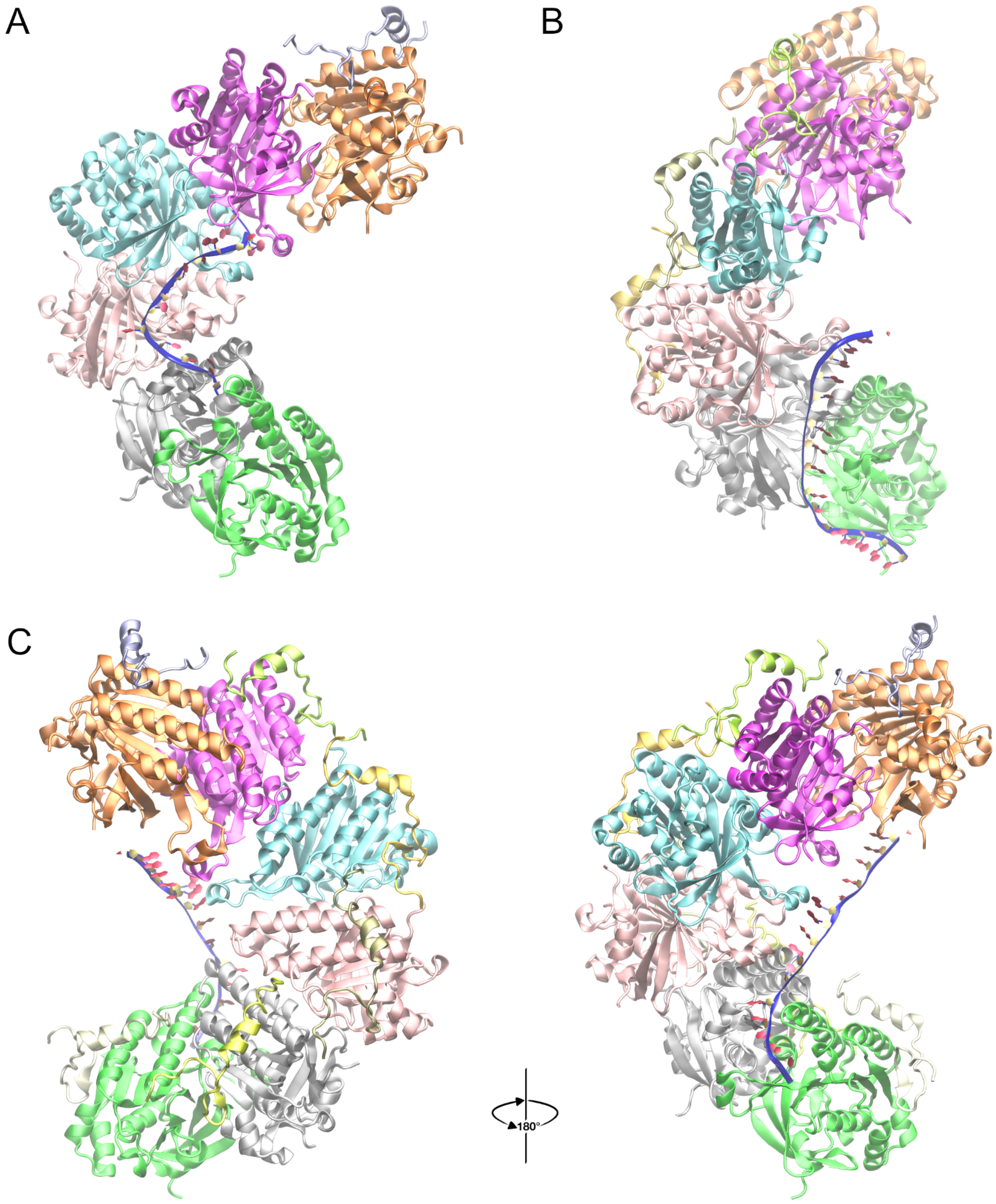


**Supplementary Figure 6. Structural prediction of RAD51 fibrils/oligomers in the presence of both the BRC4 peptide and Apt1. A**: Single RAD51 filament (6 protomers), displaying the characteristic helical structure in complex with the BRC4 peptide and Apt1 at a ratio of 1:1:1. In this model the peptide is bound to chain A (in orange), at the same spot and with a similar binding mode as in Pellegrini *et al.* (PDB ID: 1N0W)[1], whereas the Apt1 binding is occurring in the same groove as in **Supplementary Fig. 2B** (also **Fig. 3B**), but in a shifted position along the RAD51 protomers with respect to that pose: interacting with chain B, C, D, E, F, and not with chain A (in orange); **B**: RAD51 fibril in complex with BRC4 peptide and Apt1 at a 1:3:1 ratio; **C**: RAD51 fibril in complex with BRC4 peptide and Apt1 at a 1:6:1 ratio (together with its 180° rotated view, right). In the presence of six BRC4 copies (each interacting with one protomer), the only Apt1 copy, even if still interacting with the filament, loses both its bound-conformation (shown in **Supplementary Fig. 2B**) and most of the specific contacts with the RAD51 protomers. In this case, binding is mainly achieved by means of non-specific interactions via backbone.

Each protein chain is shown in a different color using the cartoon-based representation. The only aptamer copy is represented as a blue ribbon, with sugar rings in yellow and nucleobases in red. The BRC4 peptide interacting with chain A is shown in ice-blue, whereas the other BRC4 copies are shown in shades of yellow.

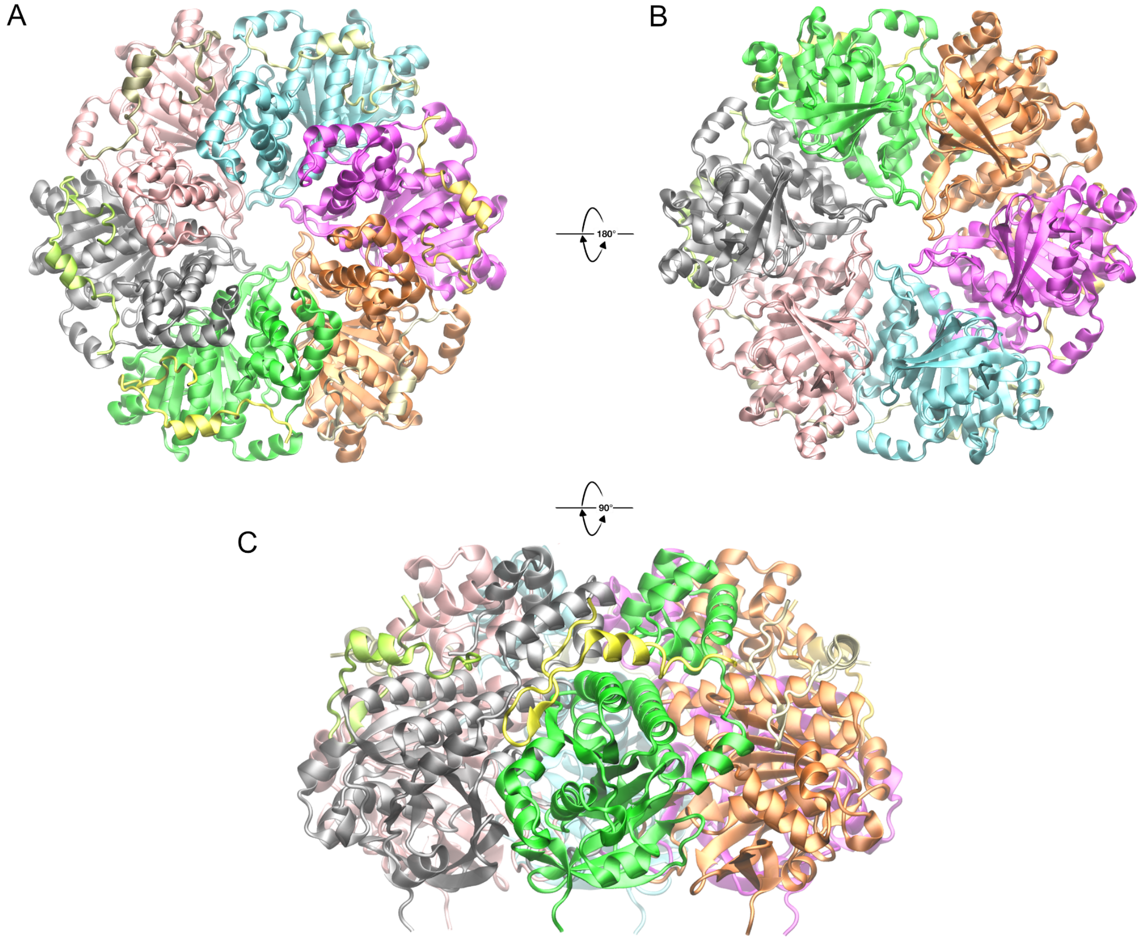


**Supplementary Figure 7. Structural prediction of RAD51 oligomer with excess of BRC4 peptide.** All the structures represent the same prediction from four different points of view of the RAD51 six protomers (not forming a helical filament structure), in complex with six BRC4 peptides (each interacting with one protomer at the experimentally determined binding spot, like in Pellegrini *et al.*[1], but in a slightly different binding conformation of both the terminals due to the presence of the N-terminal domain of the bound RAD51 protomer and the adjacent one). **A:** Top view, the resulting hexameric ring assembly resembles the archaeal Rad51 homolog structure (PDB id: 1PZN[3] and 4DC9[4], from *P. furiosus* and *M. voltae*, respectively); at the forefront the N-terminal domains together with the BRC4 peptides and their interaction patches (Region 1 and 2, including the FxxA and LFDE motifs, respectively); **B:** Bottom view (+180°), it can be appreciated how the Apt1 binding site is overlapping with the inter-chain interface at the center of the ring, thus not allowing, in this particular tight conformation, any potential interaction with the DNA aptamer; **C:** Side view (+90° tilted with respect to A, and -90° to B) centered on the F protomer showing its interaction with BRC4 (yellow), in particular the conformation of the terminals.
